# Shifts in demography in changing ecological conditions in a dependent-lineage population of harvester ant colonies

**DOI:** 10.64898/2026.03.06.710041

**Authors:** Felix Glinka, Erik B. Steiner, Eyal Privman, Deborah Gordon

## Abstract

Environmental stress can influence population dynamics and shift selection pressures. We considered the demographic effects since 2011 of intensifying drought in a J1/J2 dependent-lineage population, monitored since 1988, of the red harvester ant, *Pogonomyrmex barbatus.* A queen must mate with males of both dependent lineages to found a viable colony; mating with the same lineage results in a gyne, while mating with the other lineage results in a worker.

We used reduced representation genomic sequencing of workers from 407 colonies, ages 1-3, mostly sampled in 2021-23, to determine colony lineage, identify sets of colonies with the same mitotype, and estimate kinship between mother-daughter pairs. We found that the rare lineage J1 became even more rare, from 39% in 2011 to 25% in 2023. This creates pressure on J1 to produce more sperm or more males to allow the population to persist. We compared survival and numbers of offspring produced in the two lineages and in sets of colonies with the same mitotypes, using census data. Survival and fecundity were similar among 5 J1 and 6 J2 mitotype groups, showing no evidence for selection favoring any mitotype. We identified 62 mother-daughter colony pairs. These included 14 grandmother-triplets, with most grandmothers found in J2, suggesting that some families dominate each mitotype group. Within the 62 mother-daughter pairs, those from J1 tended to live longer than J2 colonies, maintaining a low but constant fecundity rate, while J2 colonies showed the highest fecundity at ages 11-17. Our results suggest that increasingly harsh ecological conditions are shaping population dynamics more rapidly than selection for phenotypic traits that could facilitate adaptation.

## Introduction

A fundamental question about biotic response to climate change is whether demographic decline will be slow enough to allow for adaptation by natural selection (Bell, 2013). One important constraint on the rate of adaptation is generation time, while accelerating climate change is rapidly transforming environmental conditions. Climate change is intensifying arid conditions due to rising temperatures and reduced rainfall in many parts of the world (Hatin et al., 2019). Life history strategies reflect selection pressures created by environmental and physical constraints (Healy et al., 2019), including rising temperature and water stress (Costa, 1995; Holtof et al., 2021; Manlick et al., 2021; Padda et al., 2021; Weaver et al., 2024). There may be a trade-off between survival and fecundity; reproductive allocation depends on the costs of reproduction and of survival as organisms age (Harris & Uller, 2009; Kirkwood & Rose, 1991; Lemaître & Gaillard, 2017; Stearns, 2000).

We draw on a long-term study of a population of the red harvester ant *Pogonomyrmex barbatus*, censused since 1988 (Sundaram et al., 2022). Colonies can live for 30 or more years (Gordon & Kulig, 1998), founded by a single queen who produces all of the workers, as well as reproductive daughters and sons (Suni et al., 2007). *P. barbatus* reproduces in an annual mating aggregation of the reproductives from throughout the population of colonies. Newly mated queens found new colonies that live for 20-30 years (Sundaram et al., 2022). Thus, colonies are the reproductive individuals.

An intensifying drought and decreased rainfall since about 200, across the southwestern US (Williams et al., 2020), has decreased the food supply for this species. The ants eat the seeds of annual plants and grasses that are mostly distributed from the area around the site by wind and flooding (Gordon, 1993). Regional declines in rainfall limit seed production. An increasingly limited food supply has led to increased intraspecific competition for foraging area, so that crowded colonies and founding colonies are less likely to survive in years of low rainfall, leading to an overall decrease in population size (Sundaram et al., 2022), Fig. 1).

**Figure 1.**
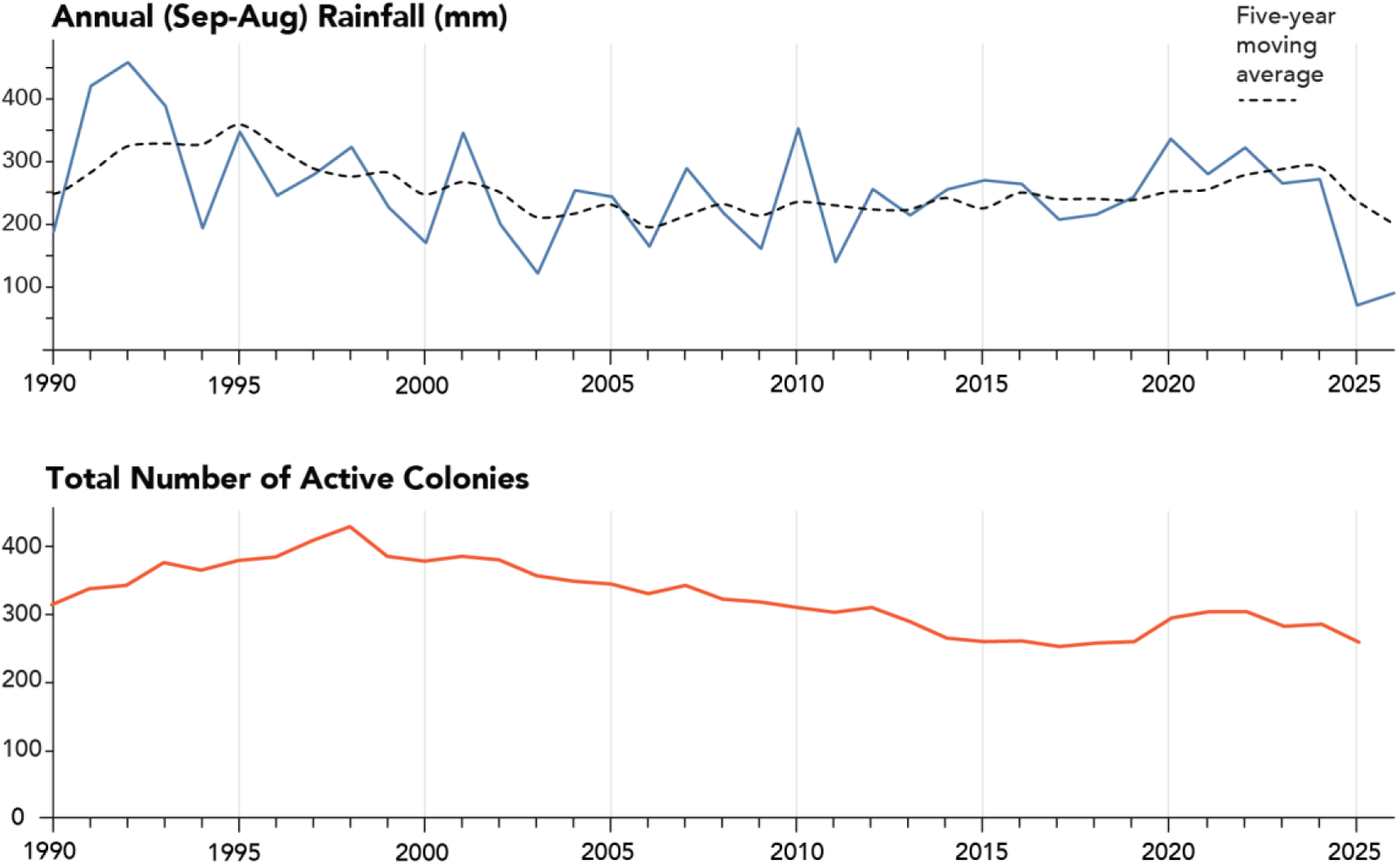
Annual rainfall 1990-2025 and total number of active colonies at the long-term data site. Rainfall data are from a weather station in San Simon, using data from a weather station about 65 km from the study site.

We considered the effects of drought conditions on the population dynamics of the unusual dependent lineage system in P. barbatus and the closely related species *P. rugosus* (Helms Cahan & Keller, 2003; Volny & Gordon, 2002). As in all Hymenoptera, males are haploid, and females are diploid. The diploid daughters of matings between a female and a male of different lineages are workers; the diploid daughters of matings between a female and a male of the same lineage are reproductives. Each queen must mate with males of both lineages to produce a viable colony. Previous work showed that queens mate with multiple males, with an effective mating frequency with the other lineage of about 4 (Volny & Gordon, 2002).

To investigate changes in the demography of the two lineages since previous studies through 2011 (Gordon et al. 2013; Ingram et al. 2013), we conducted reduced representation genomic sequencing of at least two worker samples from each of 407 colonies, ages 1-3, mostly sampled in 2021-23. We first considered changes in the lineage ratio as the drought intensified since 2011. J1 tends to be rarer than J2 in all dependent lineage populations of *P. barbatus* (Anderson et al., 2009; Gordon et al., 2013), although theory suggests the ratio should be equal (Boomsma & Grafen, 1991; Seger & Stubblefield, 2002). We next considered whether the lineages differ in fecundity, mortality, or the age of reproduction. In most animals, there is a tradeoff between life expectancy and reproduction (Blacher et al., 2017; Vergara-Martínez et al., 2024). There is no evidence for reproductive senescence in this population (Ingram et al., 2013). The lineages may differ in the age distribution of reproduction throughout the colony’s lifetime. Previous work up until 2011, at the beginning of the drought, showed no ecological advantage to either lineage (Gordon et al., 2013).

We asked whether the lineages differ in the extent to which certain families dominate the population. We considered natural selection, including survival and production of offspring colonies, by comparing mitotype groups. Because the mitochondrial haplotype (mitotype) is maternally inherited, mother-daughter pairs share a mitotype, though unrelated colonies can also have the same mitotype. If strong selection is favoring some heritable phenotypic trait that varies among mitotypes, then some mitotype groups would show higher fecundity or lower mortality than others. Mitotype groups reflect natural selection in fish (Bernatchez, 2001; Brunner et al., 2001) and elephants (Eggert et al., 2008). In ants (*Formica rufa* group, *Formica exsecta*), divergent mitotype groups in different locations indicate evolutionary pressures that may have led to recent speciation (Goropashnaya et al., 2004, 2007). By contrast, similar survival and production of offspring colonies among mitotype groups in the harvester ant population would suggest that ecological conditions affecting all phenotypes are currently shaping population dynamics.

To compare fecundity and life history in the two lineages, we identified mother-daughter pairs of colonies. We used population genomic sequencing data to find single-nucleotide polymorphism (SNP) loci and compile a genome-wide map of genetic polymorphism (Flanagan & Jones, 2019). The unusual population genetics of the dependent lineage system challenges standard methods. To address these challenges, we developed a kinship inference method appropriate for the dependent-lineage system, using lineage-specific alleles to identify mother-daughter relationships.

## Methods

### Census

Since 1988, more than 1200 colonies have been monitored at a long-term study site near Rodeo, New Mexico, USA, so the ages and locations of all colonies are known (Sundaram et al., 2022). Our genetic analysis of lineage, mitotype group, and kinship used samples of workers from 407 colonies of known age, which were founded between 1983 and 2022, collected at the study site beginning in 2000. We analyzed colony fecundity through 2023 and colony mortality through 2024.

Colonies usually begin to reproduce at age 5, with a few colonies that reproduce at age 4 (Gordon, 1995). Here, we assumed that a colony could be a mother at the age of 4. Our sample included at least 40% of the potential parent colonies in the population in every year between 2012 and 2022 (Fig. 2A&C), and 70% of the offspring colonies, counted as those found at age 1 year (Fig. 2B&C). The oldest colony sampled (in 2020) was 36 years old at the time. A higher proportion of the population was sampled in 2018-21 than in earlier years (Fig. 2C).

**Figure 2:**
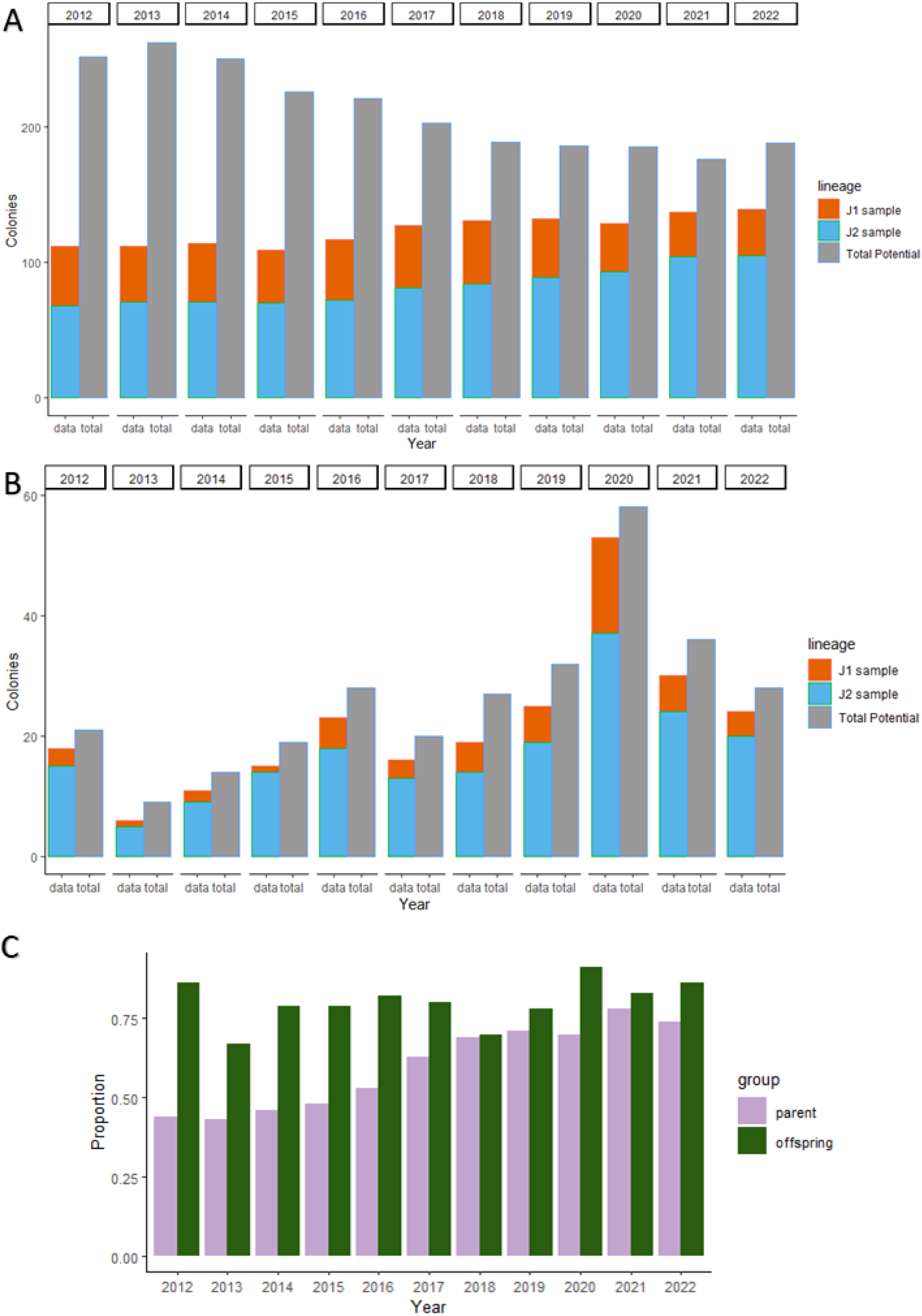
Proportion of colonies in the population that were sampled. A: Number of potential parent colonies in sample by lineage (orange and cyan) and total number of potential parent colonies (grey) per year; B: Number of newborn colonies by lineage in sample and total number of offspring colonies. Orange J1; cyan J2, grey total. C: Proportion of total potential parent colonies (blue) and offspring (green) that were sampled.

### Mitochondrial sequencing and lineage identification

Universal insect *cox*1 primers modified for *P. barbatus* (forward C1-j-1751 (Pb) C1-N-2191 (Pb) 5′) were used to amplify a 433bp portion of the mitochondrial gene *cox*1, as in Cahan & Keller (2003). All reactions were run in a 25 μl volume using Q5 Hot Start Polymerase NEB with the following PCR conditions: 98 °C for 30 seconds, followed by 35 cycles of 98 °C for 10 s, 56 °C for 30 s, and 72 °C for 30 s, paused and completed by 72 °C for 2 min. PCR products were cleaned using Ampure XP magnetic beads and sent off for Sanger sequencing (Hy Laboratories Ltd). Resulting mitotypes were used to distinguish between the two lineages.

Mitotypes were aligned with *MAFFT* (Yamada et al., 2016) and viewed in *JalView* (Waterhouse et al., 2009). All sequences were trimmed, so each sequence was of comparable size and had no low-PHRED quality base pairs at its edges. After that, the maximum likelihood gene tree was built using *RaxML* version 8 using the GTRGAMMA substitution model (Stamatakis, 2014), which resulted in two clearly distinct monophyletic clades. The lineages were then labelled as J1 and J2 based on previously defined J1 or J2 sequences in each of the clades.

### ddRAD sequencing

Double-digest Restriction site-Associated DNA sequencing (ddRAD-seq) was conducted for 1101 workers from 407 colonies (2-4 workers per colony), using a modified protocol, according to Brelsford et al. (2016), which was based on Parchman et al. (2012). Briefly, genomic DNA was extracted from the head and thorax of individual worker samples using the QIAGEN DNeasy protocol. DNA was digested using the restriction enzymes EcoRI and BfaI, which were chosen for high sequencing efficiency, as estimated using *ddgRADer* (Lajmi et al., 2023). Adaptors containing unique barcodes were ligated to each sample for multiplexing. Ligated products were PCR amplified in replicates of four using Q5 Hot Start Polymerase NEB for 20 cycles with a starting DNA volume of 6 μL. Samples were pooled and size-selected for fragments of a size of 300-700bp using BluePippin (Sage Science). The libraries were sequenced on an Illumina HiSeq X sequencer, using a paired-end, 150bp read protocol. On average, 1,519,508 reads per sample were obtained.

### ddRAD sequence data processing

The resulting reads were quality-checked with *fastQC* v. 0.12.1 (Andrews, 2010). Reads were demultiplexed using the *process_radtags* of the *STACKS2* (v. 2.66) pipeline (Catchen et al., 2011) by their barcodes and then trimmed using the program *PEAR* (Zhang et al., 2014). The resulting reads were mapped to the *P. barbatus* reference genome (J1 genome, Glinka et al. (2026)) using *bowtie2* (v. 2.5.3; using the –end-to-end and –very-sensitive parameters) (Langmead & Salzberg, 2012). Then the *STACKS2* pipeline (Catchen et al., 2011) was applied to identify SNPs and call genotypes of each sample in each locus. First, ref_map.pl from *STACKS2* was used to call SNPs, and the *populations* program was used to convert the calls into readable VCF files (using the --vcf parameter), one VCF file containing SNPs and another containing read haplotypes (where a locus corresponds to the concatenation of all SNPs within the same pair of RAD-seq reads). The read haplotype VCF has an allelic richness of 5.8 Alleles per locus, allowing for more unique alleles for each lineage. Custom scripts (https://doi.org/10.5281/zenodo.17459734) were predominantly used to process further VCF files. The SNP VCF file was used to add the depths to the respective variants of the read-haplotype VCF. We used *VCFtools* v. 0.1.13 (Danecek et al., 2011) to filter the resulting read-haplotype VCF file. We first filtered samples for missing data of over 30% (--missing-indv, then --remove), then to limit the number of missing genotypes per locus to 25% (--max-missing 0.75), requiring at least 1% minor allele frequency (--maf 0.01) per locus, and a minimum depth of 10 reads covering a locus to call the genotype of a sample (--min-DP 10). In total, we achieved a SNP depth of 76.2823 for every SNP.

### Lineage-specific alleles

Because inter-lineage matings result in sterile workers, there is no gene flow between the two lineages, resulting in an unusual distribution of genotypes in the hybrid workers. This is apparent from the striking level of heterozygosity and the unusual U-shaped distribution of heterozygosity in workers (Suppl. Fig. 2). Thus, we developed a new method to infer relatedness between colonies in a dependent-lineage population. Workers receive alleles from parents of different lineages. To infer kinship between a pair of mother-daughter colonies using worker samples, we used only the genetic material from the mother’s lineage. Workers from a mother-daughter colony pair are expected to share 25% of their maternally inherited genes.

We first identified lineage-specific alleles, using haplotypes consisting of multiple SNPs found within some of the RAD-tag loci. These are covered by the two 150-base reads obtained by paired-end sequencing, which often harbor multiple SNPs. We used a custom Python script (findAllelesInVCF.py) to identify lineage-specific alleles in the worker genotype data, with additional support from whole-genome sequencing of 12 haploid male samples from each lineage (10-18X genomic coverage; Glinka et al. (2026)). An allele was considered to be lineage-specific when it was found in at least one male sample of one lineage and not in any male sample of the other lineage, and was in heterozygous genotypes in at least 5% of all the workers, with no more than 1% homozygous genotypes. This may have allowed some alleles that were not completely lineage-specific, yet were strongly lineage-biased.

The read-haplotype VCF file was divided according to the respective lineage-specific alleles: one VCF file for each lineage, containing only loci with lineage-specific alleles of the maternal lineage, with samples from colonies that were assigned to this lineage based on the mitotype. We then created a VCF file with the haploid genotypes from the maternal lineage of each sample using another custom script (makeVCFHomozygous.py) that transformed all heterozygous genotypes into homozygous genotypes for the lineage-specific allele. We left homozygous genotypes unchanged.

### Kinship inference

To identify mother-daughter pairs of colonies, we used lineage-specific alleles of the queen’s lineage to infer kinship between two workers from a mother-daughter colony pair. A queen is the daughter of a queen and a father of the same lineage, while a worker is the daughter of a father from the other lineage. Thus, a worker from the mother colony and a worker from the daughter colony share genetic material only through the maternal lineage.

We tested for relatedness between all colonies that were potential mother-daughter pairs. As daughter colonies must have the same mitotype as their mothers, mother-daughter colony pairs could occur only within mitotype groups. We tested all colony pairs with an age difference of at least 4 years so that one could plausibly be the mother of the younger one (Gordon, 1995). We identified 11 mitotype groups in all 407 colonies, 6 for the J2 lineage and 5 for the J1 lineage.

We had 2 workers from most colonies in our sample, so there were 4 relatedness tests between most colony pairs. We used *KING* (Manichaikul et al., 2010) from the *SNPrelate* R package (Zheng et al., 2012, 2017) with the “King-homo” estimator (for homogenous populations) to estimate the kinship coefficient between pairs of workers based on the lineage-specific haploid VCF file.

We calibrated our method using the 490 pairs of nestmates in our sample as positive controls, which are expected to be either full- or half-sisters. We applied the method to all pairs of nestmate and non-nestmate samples from colonies of each lineage separately. This method produced mostly negative values, even though kinship coefficients should be between 0 and 1. Nevertheless, both lineages showed distinct, although partially overlapping, distributions of kinship coefficient values for nestmates and non-nestmates (Fig. 3). The median kinship coefficients in J1 are −0.147 and −0.472 for nestmates and non-nestmates, respectively. The two kinship distributions were significantly different for J1 (Kolmogorov-Smirnov test, D=0.65, p < 2.2e-16). Similarly, the median kinship coefficients in J2 are −0.187 and −0.470 for nestmates and non-nestmates, respectively, and the two kinship distributions were significantly different (Kolmogorov-Smirnov test, D=0.6, p < 2.2e-16). This confirmed that the method detects the statistical signal of relatedness between nestmate pairs relative to the background distribution of non-nestmate pairs.

**Figure 3:**
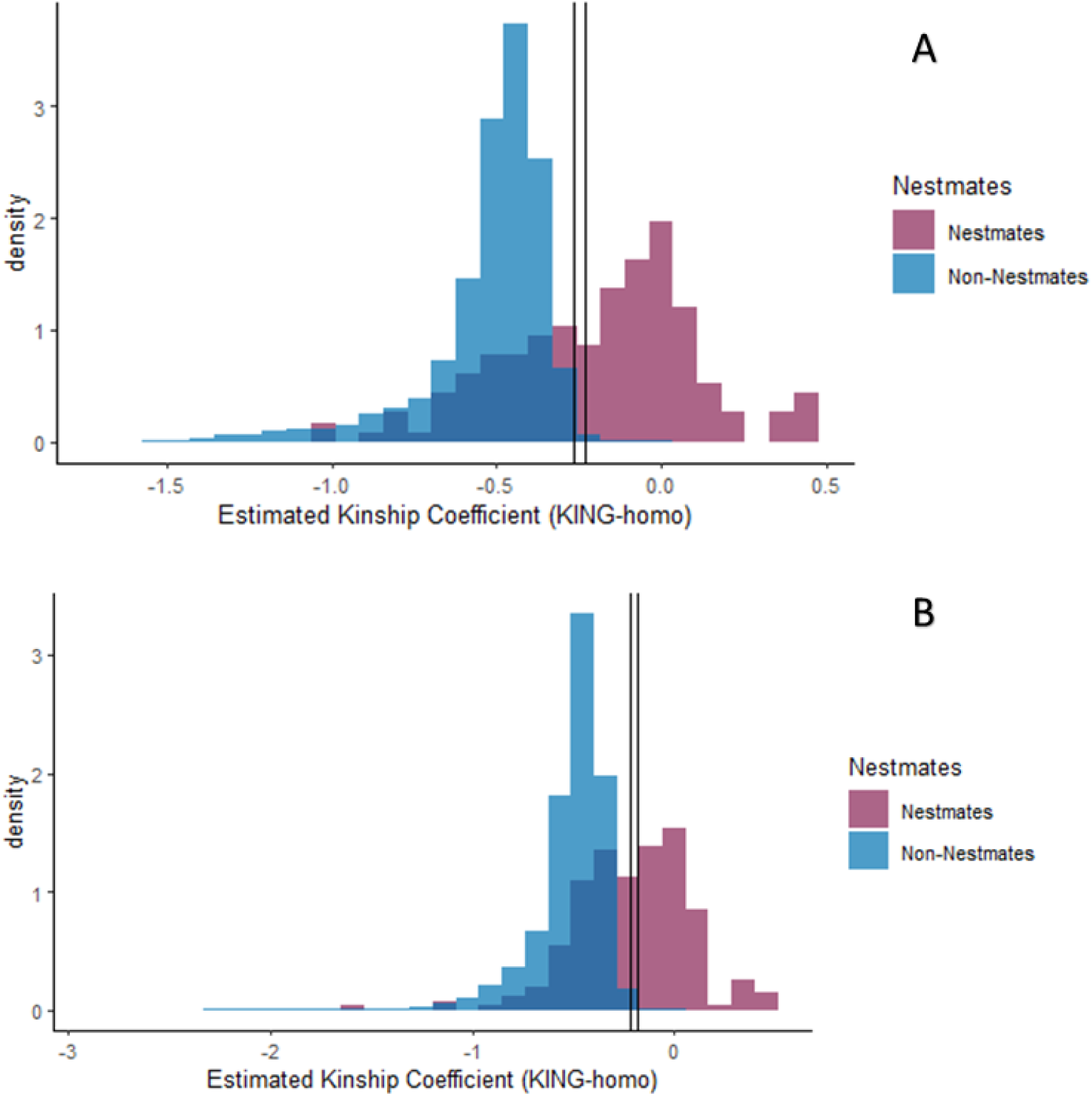
Application of the kinship inference method with nestmates as positive controls. The purple distribution shows the estimated kinship coefficient for nestmates, and the blue distribution shows the estimated kinship coefficient for non-nestmates. A. J1, B. J2.

We set a first threshold for kinship inference at −0.233 for J1 and −0.179 for J2, based on the kinship distributions of nestmates and non-nestmates. We then found all pairs of workers from two colonies of the same mitotype group that could be mother-daughter pairs (an age difference of at least 4 years) above the threshold value of the kinship coefficient. We then allowed a less stringent, slightly lower threshold of −0.266 for J1 and −0.219 for J2, for which we require at least two sample pairs between two colonies to be above the threshold to consider them a mother-daughter colony pair. Because we chose thresholds to minimize false positives, our results do not include all the mother-daughter pairs, as can be seen from the proportion of nestmates below the thresholds in Fig. 3. We calculated the true positive rate (TPR) and false positive rate (FPR) per lineage by counting the number of nestmate and non-nestmate pairs with a kinship coefficient above the threshold. For J1, with the first threshold of −0.233, the TPR is 0.6, and the FPR is 0.003. For J2, with the first threshold of −0.179, the TPR is 0.5, and the FPR is 0.001.

### Lineage ratio

We first considered whether the lineage ratio changed between 2011 and 2023. Using the data from the annual census, we examined differences in survival between the lineages of the 407 colonies in our sample. First, we tested whether living colonies of the two lineages differed in longevity by comparing the variances of the ages of living colonies using the F-test. Next, we compared the age of colony death between the lineages, using the Kolmogorov-Smirnov test. We then asked whether the lineages differed in the probability of reaching the upper quartile of the age distribution, 17 years. We compared survival per lineage below and above the age of 17 using Fisher’s exact test. Finally, we tested whether the lineages differ in survival using the Kaplan-Meier analysis with the *survfit*() function from the survival package (Therneau, 2024). We constructed a survival curve for each lineage, using all colonies sampled. We then tested whether the survival differed between lineages using the log-rank test.

To test whether years differed in colony mortality and whether this differed for the two lineages, we used a generalized linear model (GLM) modeling the counts of colony death as the response variable. We chose the model according to its AIC value (Tab. 1) and the following parameters as fixed effects: (1) years 2012 to 2024, chosen because in these years our sample included more than 40% of all colonies (Fig. 2); (2) the ratio of J1 to J2 colonies. We used a Poisson GLM because of its design for non-negative integer values, and the assumption that the variance of the data is a function of the mean, which is typical for count data (Cameron & Trivedi, 2012).

**Table 1:** Model specification.

| Model | Formula | df | AIC |
| --- | --- | --- | --- |
| Mortality |  |  |  |
| m0 | year*Lineage | 4 | 134.55 |
| m1 | year+Lineage | 3 | 132.76 |
| m2 | year | 2 | 136.42 |
| Fecundity |  |  |  |
| m0 | year*Lineage | 4 | 178.46 |
| m1 | year+Lineage | 3 | 177.67 |
| m2 | year | 2 | 256 |

We asked whether the two lineages differed in the numbers of offspring colonies by calculating the ratio of offspring (1-year-old) colonies per potential parent colony. For the offspring colonies of each year, a potential mother colony was a colony of the same lineage that was at least 4 years older. The ratio was the number of 1-year-old colonies divided by the number of potential mothers that year. We compared the distribution of this ratio in the two lineages from 2012-2022 (Fig. 2), using the Kolmogorov-Smirnov test.

To examine the possibility that some heritable phenotypic trait is associated with survival, we compared survival among mitotype groups. Our population sample contained 11 distinct mitotypes, 5 in J1 and 6 in J2. Not all colonies of the same mitotype group are closely related, but all closely related colonies must be in the same mitotype group. We first tested whether the distribution of the number of colonies per mitotype groups differs between lineages using the Kolmogorov-Smirnov test. Next, we compared the age distributions of the two largest mitotype groups of each lineage, using the Kolmogorov-Smirnov test. Finally, we compared the survival rates of the two largest mitotype groups of each lineage using a log-rank test to determine if the colonies of the mitotype groups have different patterns in their survival rates.

We inferred 62 mother-daughter colony pairs. We used these pairs to test for differences in fecundity between lineages and mitotype groups using Kolmogorov-Smirnov tests and F-tests.

To test whether years differed in colony fecundity and whether this differed for the two lineages, we used a GLM to model the counts of the number of colonies that were 1-year-old as the response variable and chose the final model based on its AIC value (Tab. 1). In the final model the fixed effects were: (1) year 2012-2023; (2) the ratio of colonies from J1 vs J2. We used a Poisson GLM for the same reasons described above for the mortality analysis.

## Results

### Lineage ratios

The number of colonies of J2 was much larger than that of J1. Of the 407 colonies sampled for this study, 120 were J1, 286 were J2, and 1 colony could not be assigned to a lineage. As of 2024, 155 of the colonies in our sample had died. Mortality was highest in 2022 (4 colonies in J1, 20 colonies in J2; Suppl. Fig. 1B).

The two lineages differed in the variance of the age distribution in 2023 (F-test, F_63,186_=1.47, p=0.05). In both lineages, the peak of the distribution was in the age group 3-4, but the J2 age distribution is more skewed to younger ages than J1 (Fig. 4A). The proportion of J2 increased from 61% (in 2011) to its maximum in 2022, 76%. The number of J1 colonies in our sample set peaked in 2020 with 71 colonies, while the number of J2 colonies peaked in 2022 with 209 colonies (Tab. 2). Lineages differed significantly in age distribution (Kolmogorov-Smirnov test, D=0.26, p=0.01; Fig. 4A). J2 had many more colonies younger than 13 years, while the difference in the number of colonies older than 13 years between J1 and J2 was only slightly higher for J2. In our sample, mortality was highest in the age group of 3-4 years (Fig. 4B), unlike in the overall population, where mortality is highest in older colonies (e.g., Sundaram et al., 2022). The lineages differed in the number of colonies that died in the upper quartile of colony age greater than 17 years (Fisher’s Exact Test, p=0.02). The survival curves of the two lineages were not significantly different (log-rank test, χ² = 0.1, df = 1, p=0.8, Fig. 5). The median survival time was 19 for J1 and 22 for J2 (Fig. 5). The death rate for J1 is lower than for J2 colonies for ages 1-10, then rises after age 10 and declines until after age 24. After age 24, survival rates are similar in both lineages.

**Figure 4:**
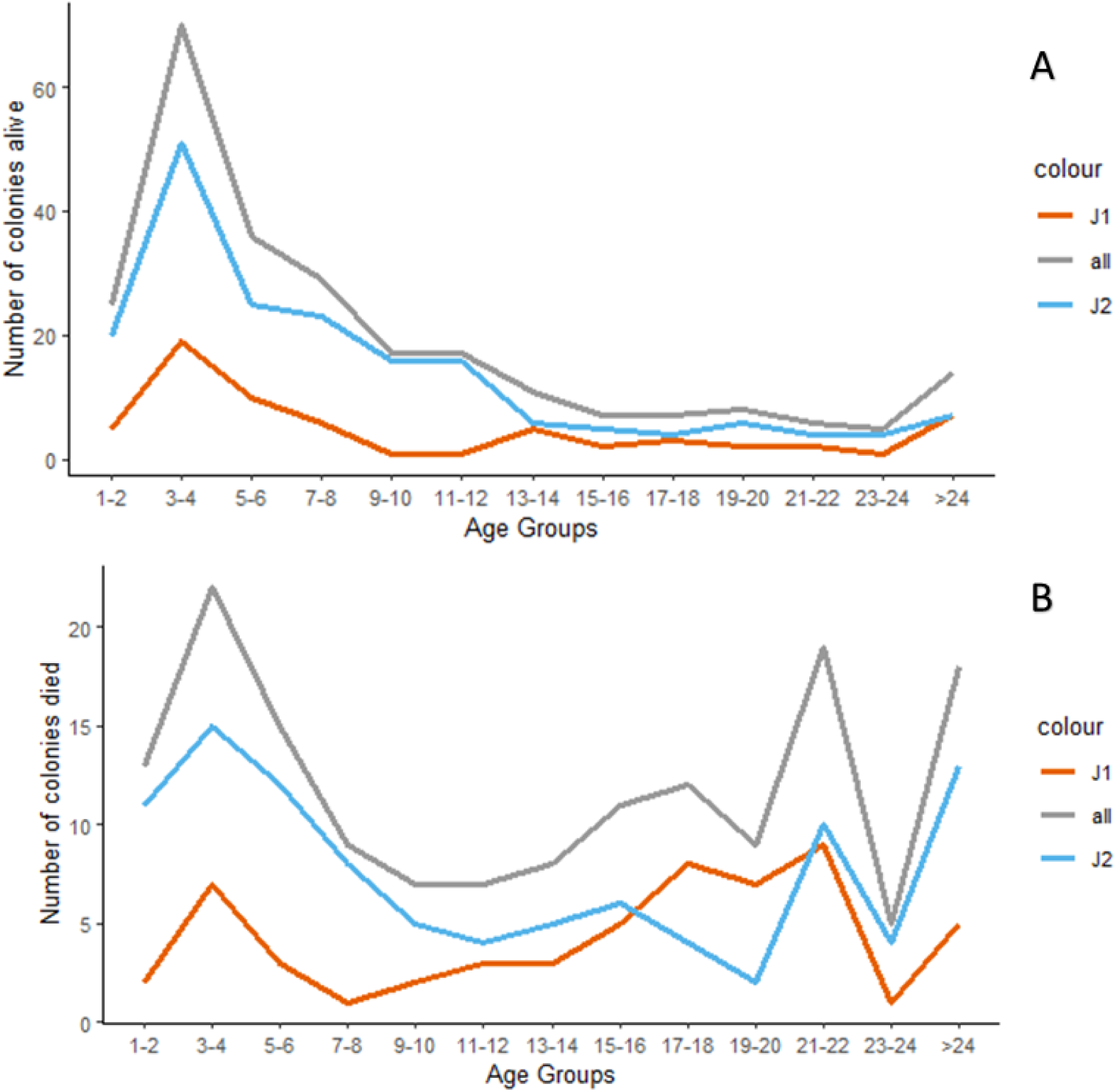
A: Age distribution of colonies in 2023 in the sample of 407 colonies. B: Distribution of age at death in the 155 sampled colonies that died. Orange J1, blue J2, and grey represent all samples together.

**Figure 5:**
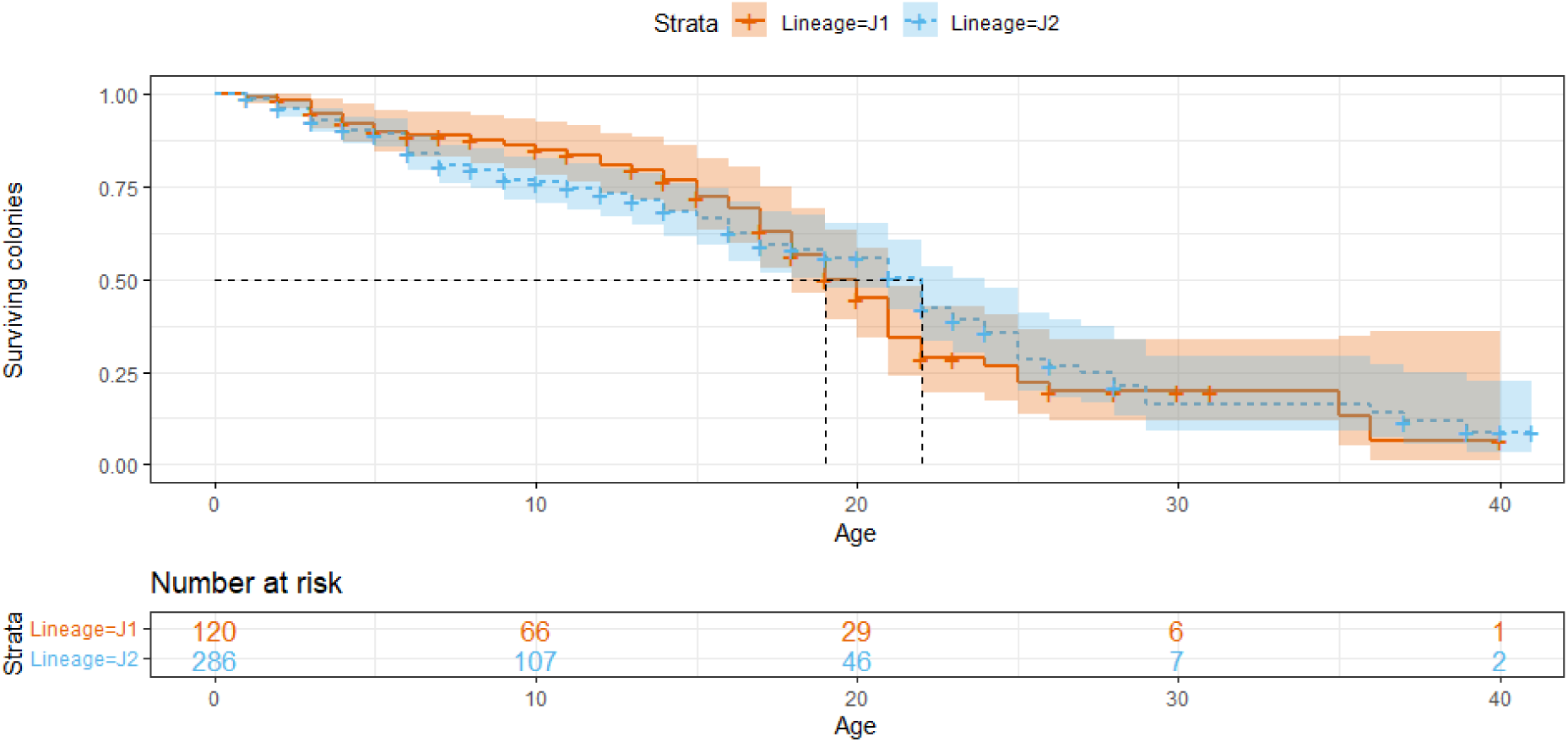
Kaplan-Meier survival plot of the two lineages. The survival probability of 50% is marked for each lineage (dotted line). The number of colonies that reached a certain age is listed in the box below. Orange J1, blue J2.

**Table 2:** History of lineage ratio of *P. barbatus* since 2011.

| Years | # J1 colonies | # J2 colonies | Proportion of J2 colonies |
| --- | --- | --- | --- |
| 2011 | 58 | 90 | 0.61 |
| 2012 | 60 | 103 | 0.63 |
| 2013 | 56 | 104 | 0.65 |
| 2014 | 57 | 108 | 0.65 |
| 2015 | 53 | 116 | 0.69 |
| 2016 | 57 | 131 | 0.70 |
| 2017 | 58 | 140 | 0.71 |
| 2018 | 63 | 151 | 0.71 |
| 2019 | 63 | 166 | 0.73 |
| 2020 | 71 | 193 | 0.73 |
| 2021 | 69 | 208 | 0.75 |
| 2022 | 68 | 209 | 0.76 |
| 2023 | 64 | 189 | 0.75 |
| total | 120 | 286 | 0.70 |

Mortality differed across years (GLM, Z=5.28, p=1.27e-07). The total number of colonies in our sample of 407 colonies still alive decreased in 2023, with a greater decrease in the J2. In J2, the largest decrease in the number of colonies over the years 2012 to 2023 was from 2022 to 2023; in J1, it was from 2020 to 2021 (Tab. 2). Before 2015, there were no differences between the lineages in the numbers of colonies that died. After 2015, mortality was initially higher in J1; after 2020, mortality was higher in J2 (Suppl. Fig. 1B; Z=2.33, p=0.02; Tab. 3).

**Table 3:** Model testing year-to-year differences between lineages in the mortality of colonies in the sample set.

|  | Estimate | Std. Error | z value | p value |
| --- | --- | --- | --- | --- |
| (Intercept) | -258.12 | 49.16 | -5.25 | 1.51e-07 |
| Year colonies died | 0.13 | 0.02 | 5.28 | 1.27e-07 |
| Lineage Ratio J1/J2 | 0.42 | 0.18 | 2.33 | 0.02 |

The two lineages differed significantly in the number of offspring colonies per year (Z=1.11, p=3.41e-16; Tab. 5), and in fecundity rate (potential offspring per possible parent) (Kolmogorov-Smirnov Test, D=0.64, p=0.02). From 2012 to 2015, the number of offspring per potential mother per year was less than 0.2 for both lineages, and was highest in 2019, with a peak of 0.31 for J1 and 0.33 for J2, and then declined to a value below 0.05 by 2022 (Fig. 6). J2 had a higher fecundity rate than J1 in the years up until 2015, but this gap between the lineages closed in 2019. In most years from 2011 to 2021, the majority of new colonies were in the J2 lineage (Fig. 2), maintaining the skewed lineage ratio (Fig. 4A).

**Figure 6:**
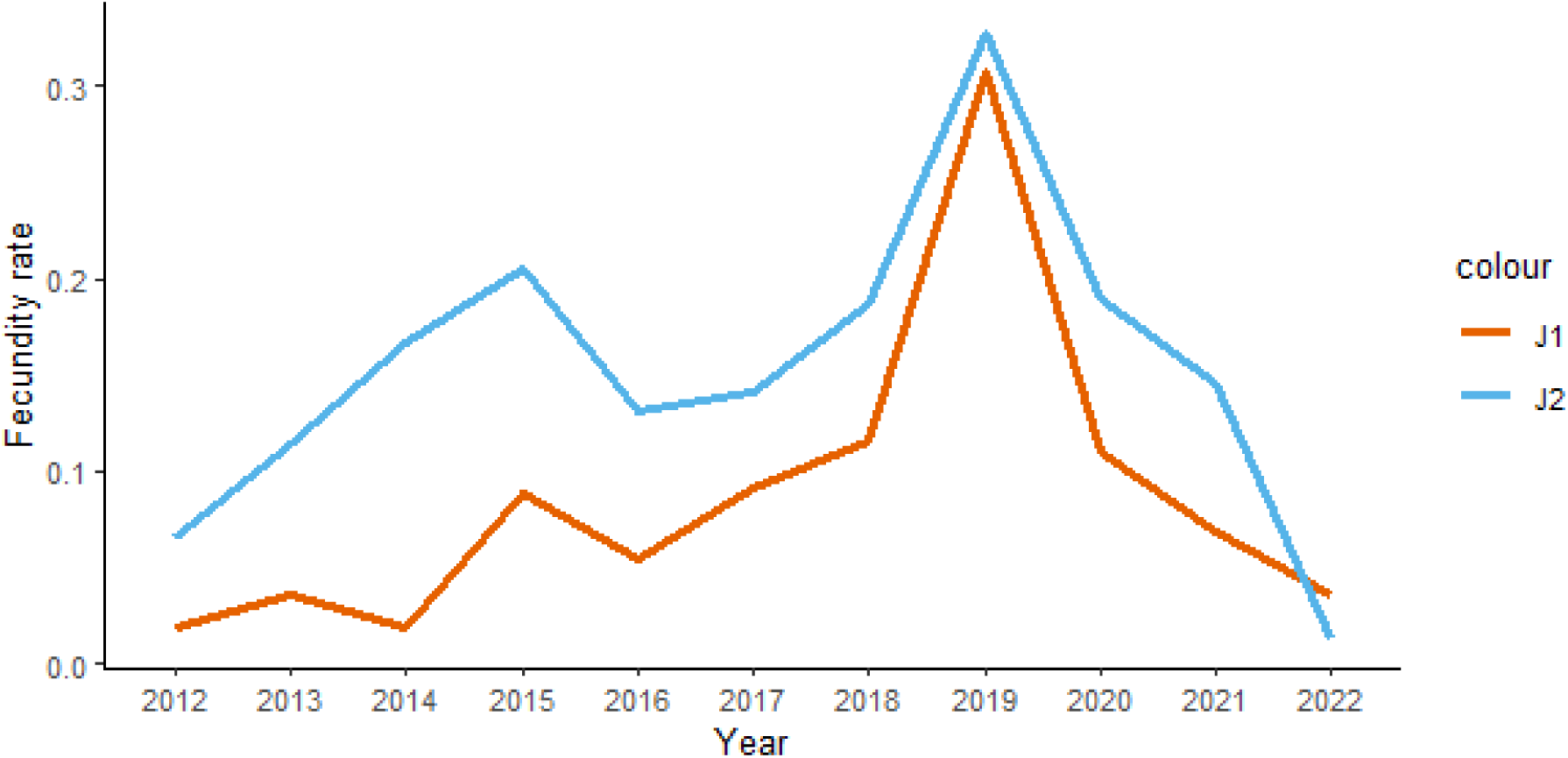
Fecundity rate (potential offspring per possible parent) for each lineage by year. Orange J1, blue J2.

### Mitotype groups

We identified five mitotype groups from 120 colonies in J1 and six mitotype groups from 286 colonies in J2 (Tab. 4; sequences in Suppl. Tab. 1). For the remaining 26 colonies, the sequences were sufficient to identify the lineage, but not a mitotype group. Both lineages have a similar number of colonies per mitotype group (Kolmogorov-Smirnov test, D=0.27, p=0.93).

**Table 4:** Mitotype groups.

| Mitotype Group | Size in the sample set |
| --- | --- |
| J1 |  |
| J1A | 82 |
| J1B | 12 |
| J1C | 11 |
| J1D | 3 |
| J1E | 2 |
| J2 |  |
| J2A | 192 |
| J2B | 50 |
| J2C | 9 |
| J2D | 9 |
| J2E | 8 |
| J2F | 2 |

**Table 5:** Model testing year-to-year differences between lineages in fecundity.

|  | Estimate | Std. Error | z value | p value |
| --- | --- | --- | --- | --- |
| (Intercept) | -117.46 | 37.9 | -3.09 | 0.002 |
| Year colonies were 1 year old | 0.06 | 0.02 | 3.14 | 0.002 |
| Lineage Ratio J1/J2 | 1.26 | 0.15 | 8.16 | 3.41e-16 |

Mitotype groups were labelled according to the number of colonies, with J1A (82 colonies, 75% of all) and J2A (192 colonies, 71% of all) the largest in each lineage, J1B (12 colonies; 11% of all J1 colonies) and J2 B (50 colonies; 19% of all J2 colonies) the second largest, and so on (Fig. 7A & B).

**Figure 7:**
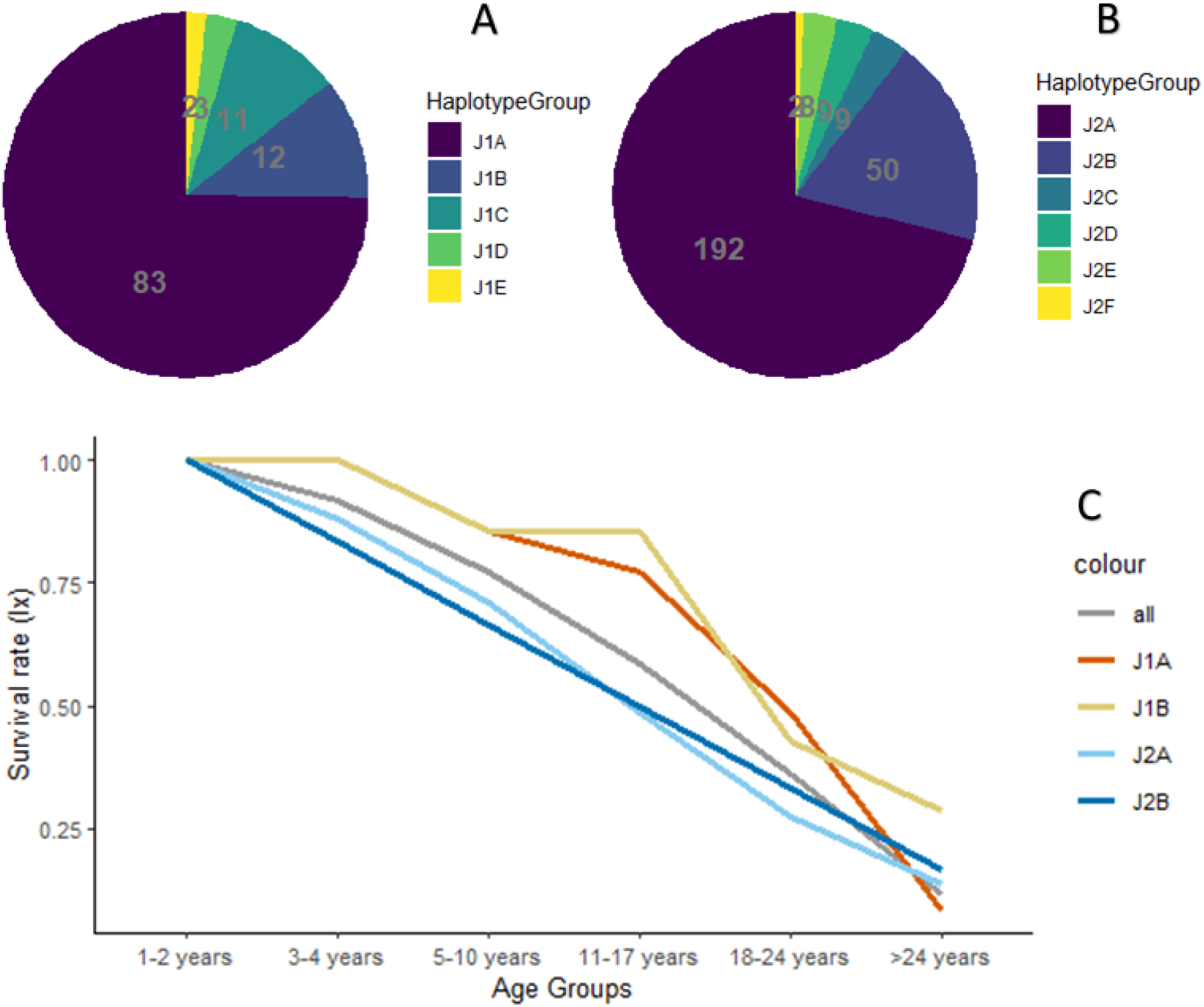
Survival plot per mitotype group per lineage. A: Mitotype groups in lineage J1. B: Mitotype groups in lineage J2. C: Survival by mitotype group for the two largest groups. J1A Light orange, J1B orange, J2A light blue, J2B blue, all samples, gray.

The two largest mitotype groups within each lineage did not differ in the age distribution of colony mortality (J1A vs J1B: Kolmogorov-Smirnov Test, D=0.29, p= 0.61; J2A vs J2B: Kolmogorov-Smirnov Test, D=0.12, p= 0.1). Within each lineage, the two largest mitotype groups of colonies did not differ in survival; for J1A and J1B (log-rank test, χ² = 0, df = 1, p=1), for J2A and J2B (log-rank test, χ² = 0.7, df = 1, p=0.4). Until the age of 17, both of the two largest mitotype groups of J1 have a higher survival rate than those of J2, up to the age of 24 (45% for J1 A and J1 B) than those of J2 (10%), as reflected in the survival rate of J1 overall (Fig. 5 & Fig. 7C)

### Mother-daughter pairs

We used reduced-representation genomic sequencing (ddRAD-seq) to identify 62 mother-daughter colony pairs based on genetic relatedness: 21 pairs of J1 and 41 pairs of J2. The 62 pairs include 14 grandmother-mother-daughter triplets, consisting of 2 of these triplets from J1 and 12 from J2, including one great-grandmother in J2. The number of mothers at reproduction peaks at ages 11-17 years (*mx* as age-specific fecundity, Fig. 8A).

**Figure 8:**
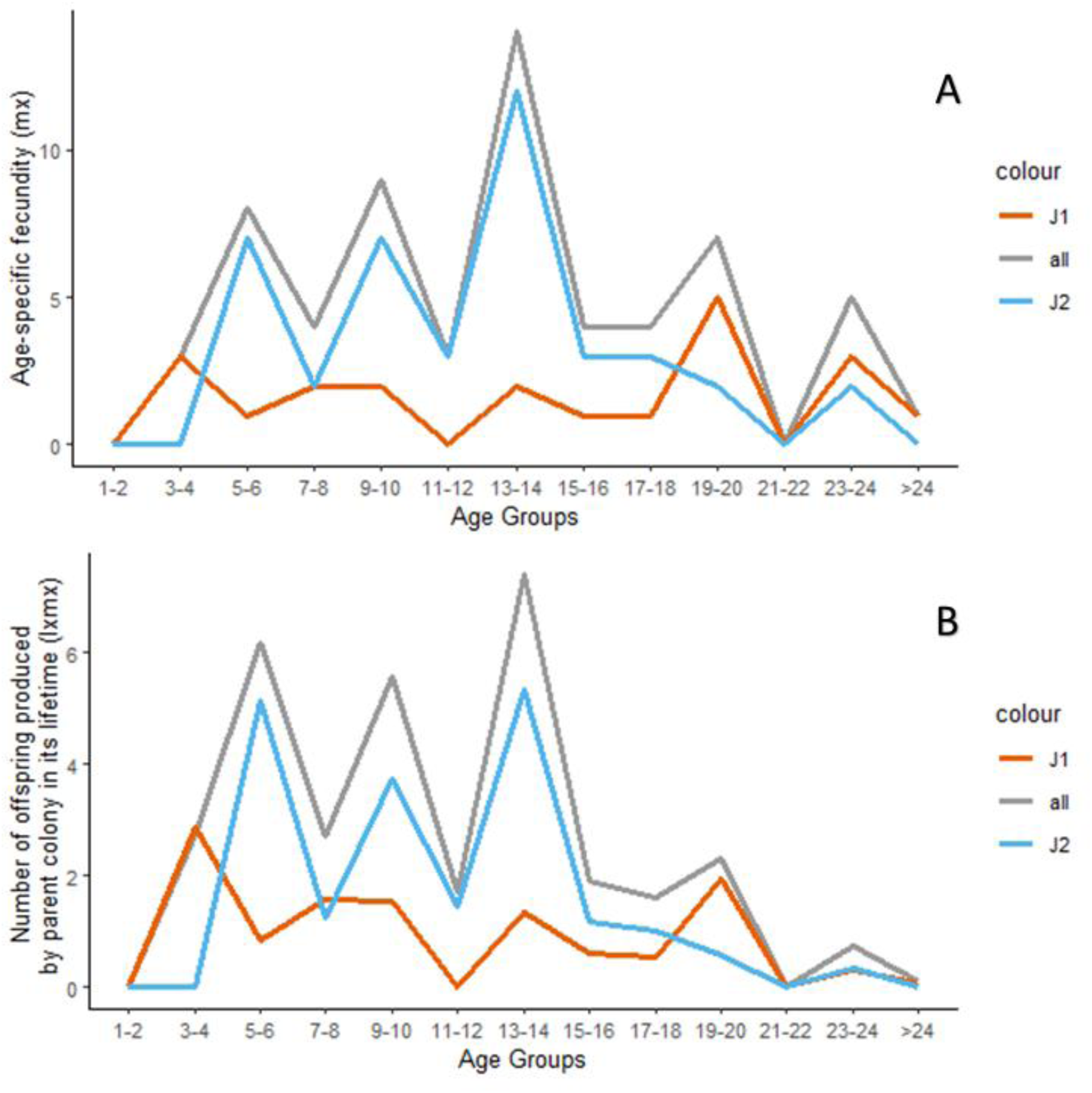
Fecundity for identified mother-daughter pairs of colonies, by lineage. A: age-specific fecundity (*mx*) per age group by lineage. B: product of age-specific survival and reproduction (*lxmx*). Orange: J1, blue: J2, and gray: all samples together.

Identified mother-daughter pairs of the two lineages differed in the distribution of age at reproduction (Kolmogorov-Smirnov test, D=0.35, p=0.02), and the variance of age at reproduction was higher in J1 than in J2 (F-Test, F_20,40_ = 2.58, p=0.01). In J1, the fecundity rate was low and steady, with a mean of 1.61 and a peak of 5 offspring in ages 19-20. In J2, the average was 3.15 offspring, with a peak of 12 offspring in ages 13-14 (Fig. 8A). Colonies became grandmothers at ages 10 to 34. The number of offspring per parent by age group for J2 peaked at 5.33 for ages 13-14 and declined to 0 offspring per mother at the ages older than 24 (*lxmx* as product of age-specific survival and reproduction (Fig. 8B).

From 2012 to 2022, we identified the parent colony for 28% of the offspring colonies (Fig. 9). This proportion differed among years, with a mean of 15% of colonies for which a parent colony was identified per year.

**Figure 9:**
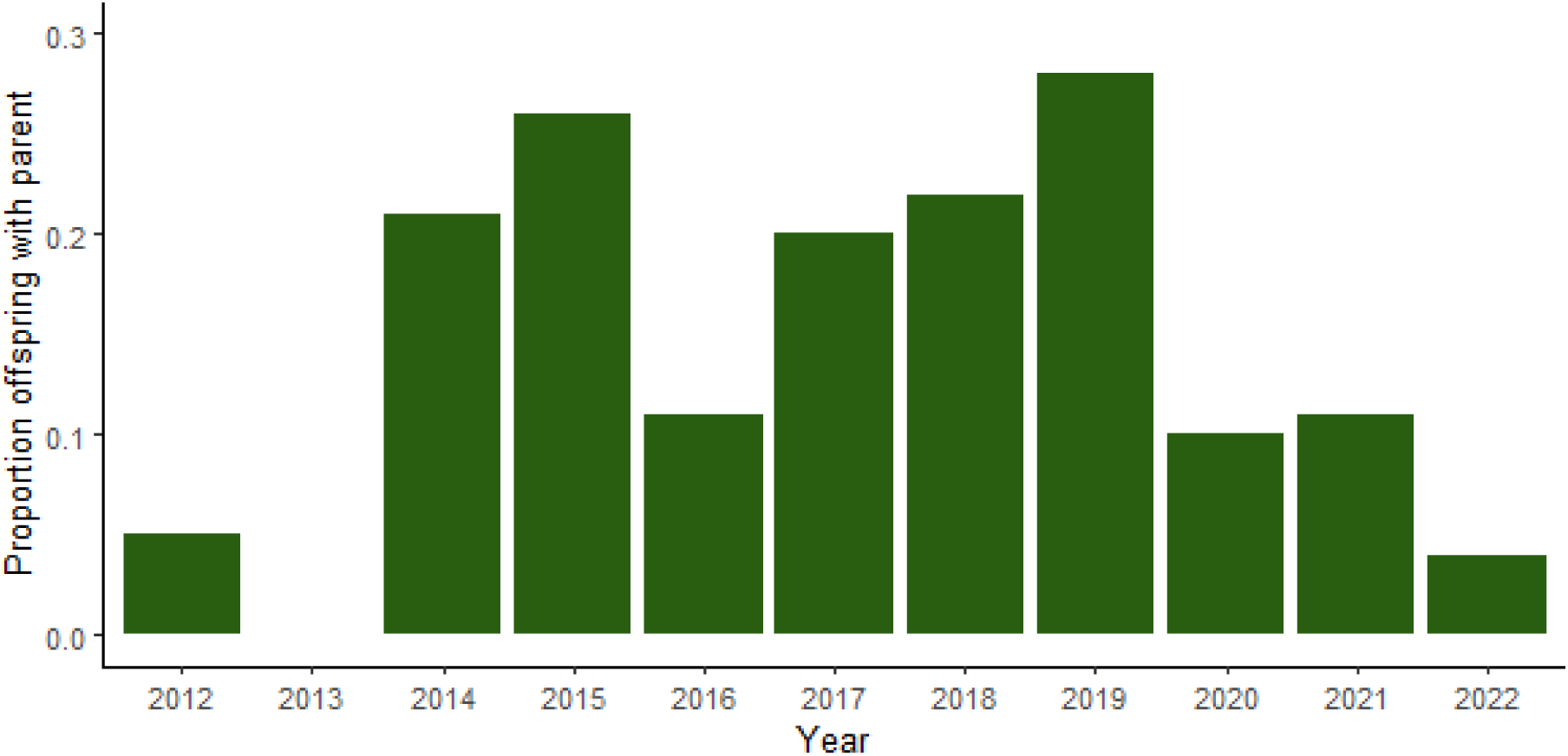
Proportion of offspring colonies for which a parent colony was identified, by year

Mitotype groups within each lineage did not differ in the age distribution of mothers that we identified (Kolmogorov-Smirnov test, D=0.25, p=1). The distribution of mothers we identified within mitotype groups was consistent with the overall numbers for each lineage (compare Fig. 7A & B, with Fig. 10A & B). Mothers we identified in the two largest mitotype groups of J1 and J2 did not differ in age at reproduction (J1A and J1B, Kolmogorov-Smirnov Test, D=0.5, p= 0.75; J2A and J2B, Kolmogorov-Smirnov Test, D=0.29, p= 0.47). Mothers we identified in J1A and J1B show low, but constant fecundity, while J2A and J2B reached their maximum fecundity at the age of 11-17 and then decreased (Fig. 10C). While the distributions of age at reproduction for mothers we identified in the mitotype groups differ slightly from the overall distributions for each lineage (compare Fig. 8A with Fig. 10C), they show the same trends: similar fecundity across ages in J1 and a peak at ages 11-17 in J2. However, the mothers we identified in J1A and J1B differ slightly, with a decline in fecundity in J1A after the age of 24 and slightly more fluctuation compared to J1B. In some years, J1 produced more offspring than J2 (Suppl. Fig. 1A; GLM Z=3.14, p=0.002).

**Figure 10:**
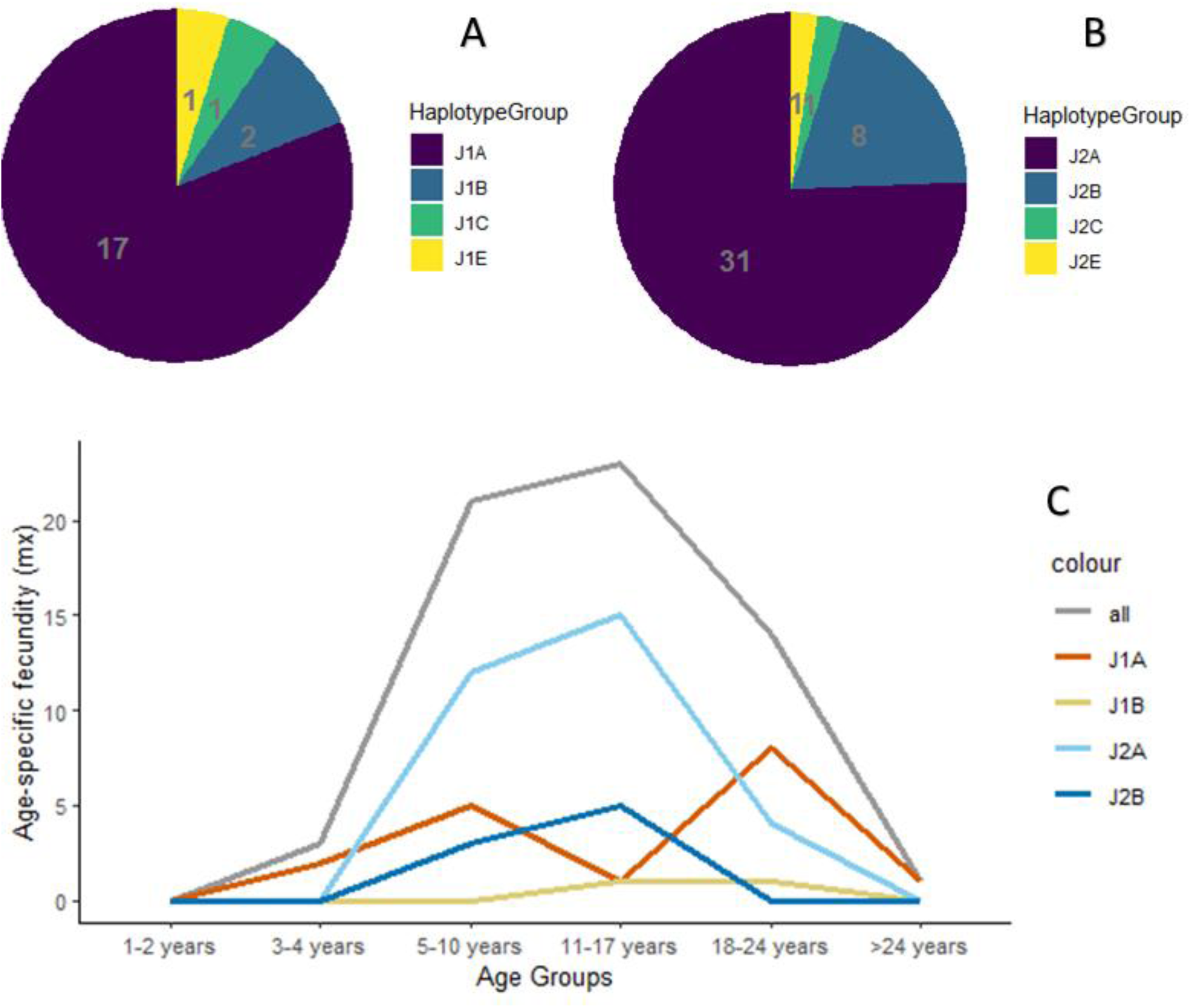
Age-specific fecundity in the mother-daughter pairs we identified in the two largest mitotype groups of each lineage. A: Distribution of the number of identified mothers per mitotype group in lineage J1. B: Distribution of the number of mothers per mitotype group in lineage J2. C: Fecundity plot by mitotype group. Light orange J1A, orange J1B, light blue J2A, blue J2B, and gray for all samples.

## Discussion

The lineage ratio was strongly skewed toward J2. The current ratio is consistent with the lineage ratio from other dependent-lineage populations of *P. barbatus* and *P. rugosus* (Anderson et al., 2009; Gordon et al., 2013; Schwander et al., 2007). The lineage ratio of J1 to J2 from 2012 to 2023 (mean: 0.7, variance: 0.002; Tab. 2) decreased about 10 % from the ratio of 64% (mean: 64.09, variance: 4.22) previously reported from 1986 to 2010 in the study population (Gordon et al., 2013).

The skewed lineage ratio raises the question of how the more common lineage, J2, can find enough mates of the rare lineage, J1. Just as sex ratios in animals are influenced by life history differences between the sexes (Kappeler et al., 2023), similar factors may help to maintain the two lineages. Queens mate with many males (Suni & Eldakar, 2011), and each male mates only once (Gordon, 2024). Previous work suggested that J1 and J2 gynes tend to mate with approximately the same number of males of the other lineage (Gordon et al. 2013). A *Pogonomyrmex* queen generally acquires sperm in the same ratio as the lineage ratio in the population (Helms Cahan & Gardner-Morse, 2013; Helms Cahan & Julian, 2010; Helms Cahan et al., 2002; Helms Cahan & Vinson, 2003).

To maintain the population with a deeply skewed lineage ratio, either J1 colonies produce more males, so that each gyne on the mating aggregation can mate with a male from the rare lineage, or each J1 male produces more sperm than each J2 male. The availability of mates of the rare lineage, J1, to gynes of J2 may be influenced by differences in the number or size of gynes produced by J2 colonies. In a study conducted in the 1990s, before the drought began, about 30% of the colonies in the population did not produce any reproductives. Older colonies tended to produce more reproductives but smaller females (Wagner & Gordon, 1999), and it is possible that smaller females are less likely to survive (Keller & Passera, 1989). Because the two lineages differ in their cuticular hydrocarbon profile (Volny et al., 2006), gynes may be able to preferentially select males from the rarer lineage. Studies on Hymenoptera show that workers may have an influence on the sex ratio within a colony (Boomsma & Grafen, 1991; Chapuisat et al., 1997; Murakami et al., 2000; Sundström et al., 1996), so they could act to increase the proportion of males in this species. Lineages may differ in the consumption of reproductive female eggs (Helms Cahan et al., 2004; Schwander et al., 2006).

Since the early 2000’s, decreasing rainfall has led to a lower food supply for this seed-eating species, and competition for foraging areas has increased. (Sundaram et al., 2022). Even before 2000, only about 1% of female alates in the population managed to found viable new colonies (Gordon & Kulig, 1998). Our results suggest that these harsher ecological conditions may create differences between the lineages in allocation to reproduction and survival (Bell, 1980; Lemaître & Gaillard, 2017). While previous work has shown no ecological differences between the lineages, there may be different pressures on life history (Kirkwood & Rose, 1991). Our results suggest a possible tradeoff between increased fecundity in J2 and survival in J1.

First, the lineages differ in age distribution, suggesting there may be an effect of drought conditions on the survival of young colonies of J2 (Fig. 5). Colonies of J1 were more likely than those of J2 to survive until age 17, but J2 colonies that survived to age 17 were likely to live longer than J1 colonies.

Overall, the number of new colonies per year has declined since drought conditions began in the early 2000’s (Sundaram et al., 2022). In the mother-daughter pairs we identified from 2012-2023, the average number of offspring per mother colony was 1.23, with a range of 1 to 3 (Ingram et al., 2013). Before 2010, the average was 2.94 with a range of 1 to 8. In our sample, fecundity was highest for mothers ages 11-17, mostly determined by reproduction by J2.

The lineages differ in fecundity. In our sample through 2023, fecundity is higher in J2 colonies until the age of 13-14 with a peak at ages 3-4, while in J1 fecundity is constant up to age 20 then drops (see Fig. 4B & Fig. 8A). This peak in fecundity differed from the results of a previous study through 2010, which showed no trends toward reproductive senescence up to age 24 (Ingram et al., 2013). Ovarian state in *P. barbatus* changes with age, suggesting a decline in reproductive potential over time (Vergara-Martínez et al., 2025). This may be more pronounced in J2 than in J1, leading to the drop in fecundity in J2 colonies after the age of 17. Despite this, J2 consistently produced more offspring than J1, increasing its proportion of the population.

The dependent lineage system introduces special challenges for the identification of kinship because it creates genotype distributions in the hybrid workers that are very different from those of most sexually reproducing populations. The reproductive isolation of two lineages has led to divergent allele frequencies and the accumulation of fixed lineage-specific alleles. The extreme and unusual distribution of heterozygosity (Suppl. Fig. 2) may distort many of the standard population genetic statistics and inference methods (e.g., Browning & Browning, 2013; Lajmi et al., 2025) because it contradicts common assumptions underlying those methods, such as random mating. Here, we developed a novel approach for population genetic analysis based on the identification of lineage-specific mitotypes from genomic sequencing data. Other studies of dependent-lineage populations may be able to apply this approach.

It appears that some families tend to dominate within each mitotype group. We identified 62 mother-daughter pairs of colonies, of which 14 (about 23%) are in grandmother-granddaughter triplets, and one great-grandmother. Similarly, in a larger sample, Ingram et al. (2013) found 19 grandmother triplets (39% of all offspring), six great-grandmothers, and one great-great-grandmother. Our results are consistent with the previous results showing that most reproduction occurs in only 25% of the potential parent colonies (Ingram et al., 2013).

However, we found no evidence for current intense selection on any heritable aspect of phenotype. Although we could identify only a small proportion of the mother-daughter pairs in the population, these were about equally distributed among mitotype groups. Since the mitotype is maternally inherited, all mother-daughter pairs must share the same mitotype. However, not all members of the same mitotype group are necessarily related, as there are only about 11 mitotypes (5 in J1, 6 in J2) in the population. If there were strong selection for a heritable phenotypic trait, we would expect to see differences among mitotype groups in recruitment or survival. Instead, each mitotype group of a lineage shows the fecundity characteristic of its lineage (Fig. 8 vs Fig. 10C), and there were no significant differences in survival (Fig. 7C).

Thus, while our study does not rule out the possibility of selection due to some heritable phenotypic trait, it shows that the intensity of such selection, if it is occurring, is much lower than the intensity of effects on survival of rapidly accelerating drought conditions. It appears that rapid effects of ecological conditions on survival may have more impact than the slower process of adaptation through natural selection.

## Supplementary

**Supplementary Table 1:**
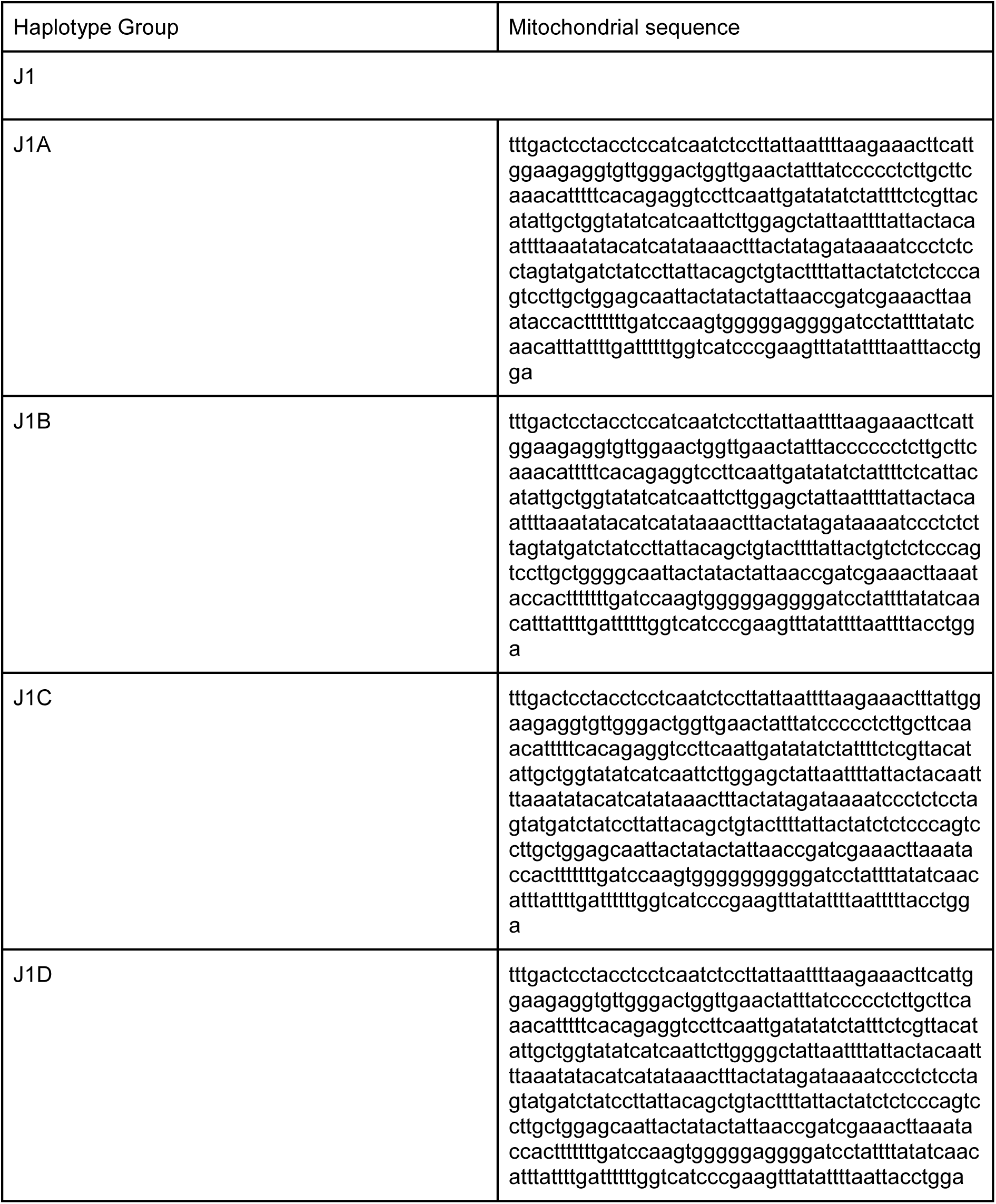
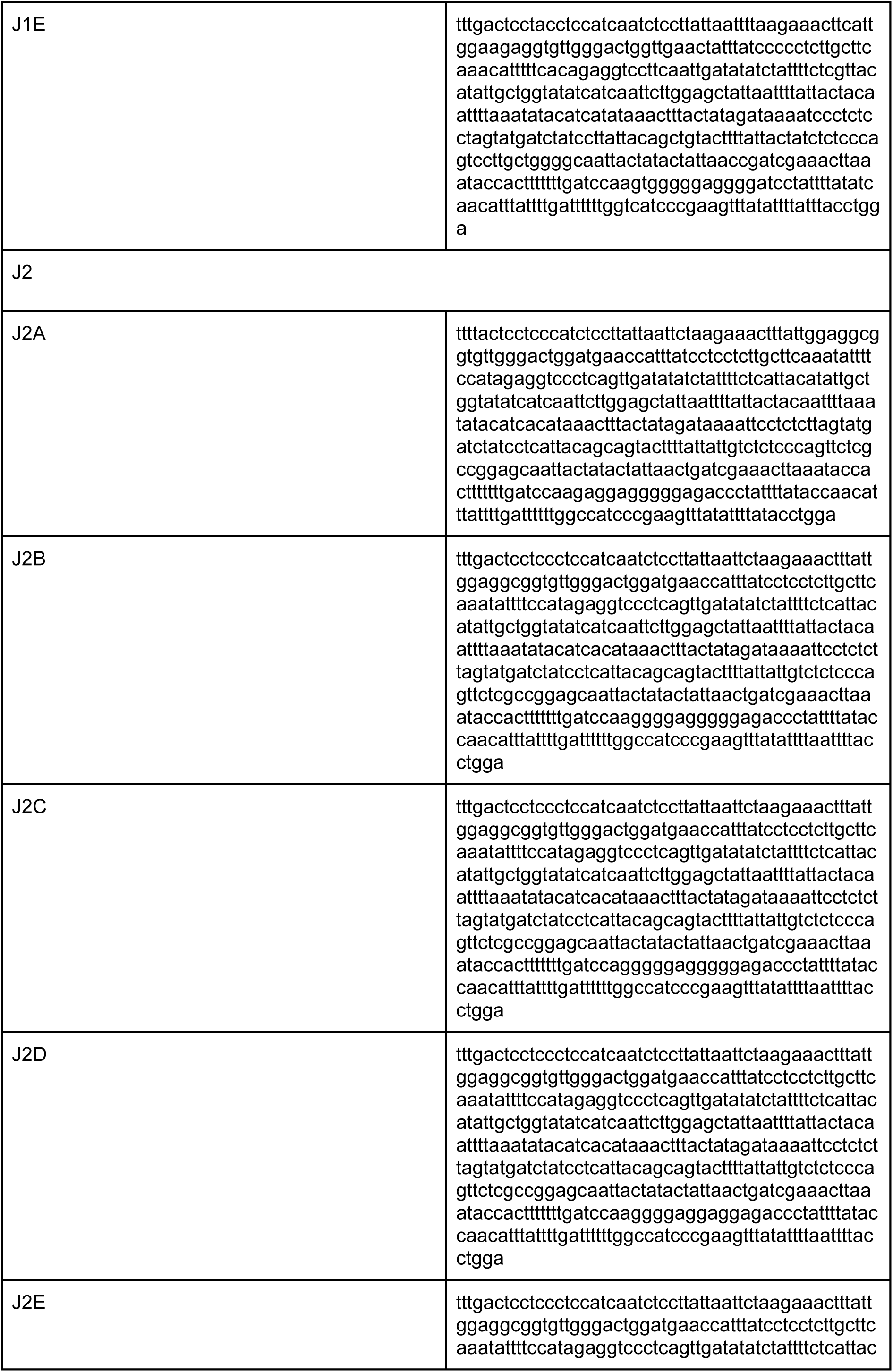
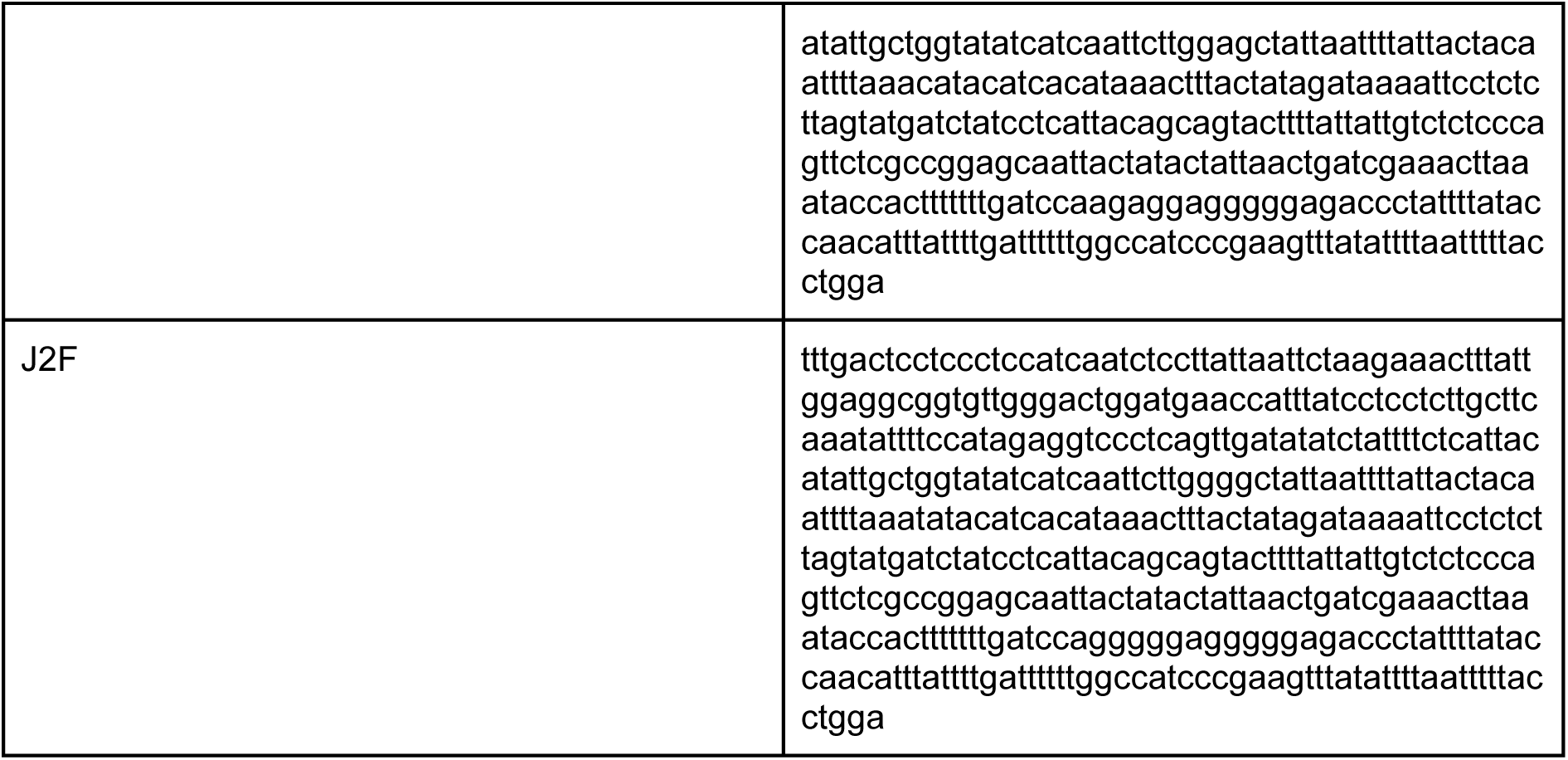
Sequences of haplotype group.

**Supplementary Figure 1:**
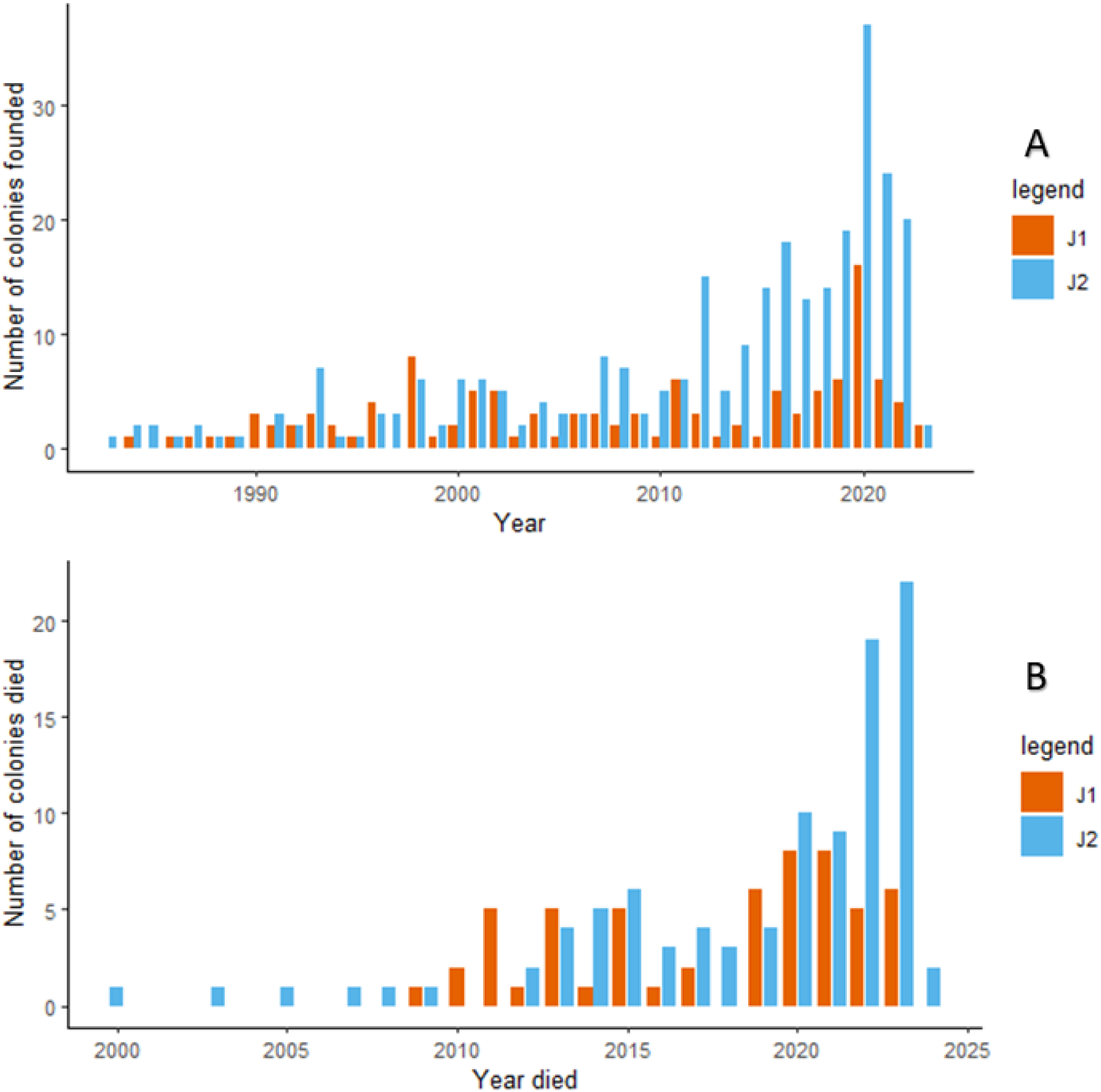
Number of colonies in the sample set found per year (A) and number of colonies in the sample set that died per year (B)

**Supplementary Figure 2:**
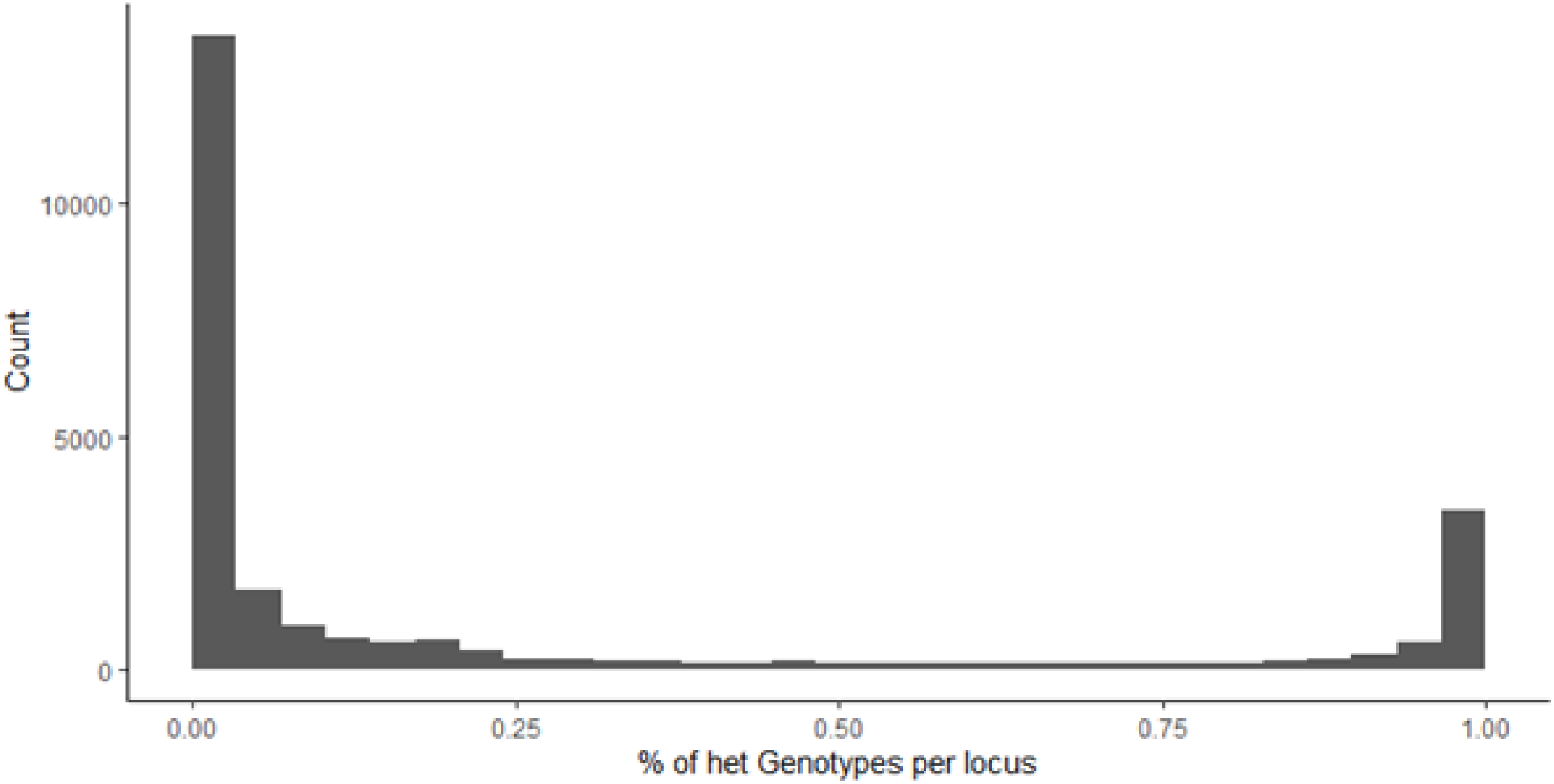
Count of heterozygous genotype per locus

## Data Accessibility Statement

The scripts to find alleles belonging to one lineage and make a vcf homozygous for one lineage used in this study were submitted to the zenodo repository (10.5281/zenodo.17459734). The ddRAD sequencing data are being processed to upload to NCBI (PRJNA1433433, PRJNA1433508, PRJNA1433511). The remaining data will be deposited in the Stanford Data Repository.

## Competing Interests Statement

The authors declare no conflicts of interest.

## Acknowledgements

The work was supported by National Science Foundation Award (1940647) to D.M.G. and U.S.-Israel Binational Science Foundation grant (2019655) to E.P. We are grateful to Aparna Lajmi and Chih-Chi Lee for guidance in the lab. Many thanks to all of the research assistants who helped with the collection of the samples over the years, especially Marco Carrillo, Henry Cerbone, Max Mazydryk, Diego Perez, Zander Opperman, Viggo Rey, and Mikaela Wilson. We thank the staff at the Southwestern Research Station for all of their help with field work. We also want to thank Hunter Fraser for valuable discussions and Antoine Melet for comments on the manuscript.

## Authors’ contributions

F.G.: methodology, investigation, formal analysis, data curation, writing – original draft, visualization

E.P.: conceptualization, methodology, resources, supervision, project administration, writing – original draft, funding acquisition

E.B.S.: data curation

D.M.G.: conceptualization, methodology, investigation, resources, supervision, project administration, writing – original draft, funding acquisition

## Notes

### Competing Interest Statement

The authors have declared no competing interest.

