## Supplementary Table 1 for "Shifts in demography in changing ecological conditions in a dependent-lineage population of harvester ant colonies"

Supplementary Table 1: Sequences of haplotype group

| Haplotype Group | Mitochondrial sequence |
| --- | --- |
| J1 | |
| J1A | tttgactcctacctccatcaatctccttattaattttaagaaacttcattggaagaggtgttgggactggttgaactatttatccccctcttgcttcaaacatttttcacagaggtccttcaattgatatatctattttctcgttacatattgctggtatatcatcaattcttggagctattaattttattactacaattttaaatatacatcatataaactttactatagataaaatccctctcctagtatgatctatccttattacagctgtacttttattactatctctcccagtccttgctggagcaattactatactattaaccgatcgaaacttaaataccacttttttgatccaagtgggggaggggatcctattttatatcaacatttattttgatttttggtcatcccgaagtttatattttaatttacctgga |
| J1B | tttgactcctacctccatcaatctccttattaattttaagaaacttcattggaagaggtgttggaactggttgaactatttacccccctcttgcttcaaacatttttcacagaggtccttcaattgatatatctattttctcattacatattgctggtatatcatcaattcttggagctattaattttattactacaattttaaatatacatcatataaactttactatagataaaatccctctcttagtatgatctatccttattacagctgtacttttattactgtctctcccagtccttgctggggcaattactatactattaaccgatcgaaacttaaataccacttttttgatccaagtgggggaggggatcctattttatatcaacatttattttgatttttggtcatcccgaagtttatattttaattttacctgga |
| J1C | tttgactcctacctcctcaatctccttattaattttaagaaactttattggaagaggtgttgggactggttgaactatttatccccctcttgcttcaaacatttttcacagaggtccttcaattgatatatctattttctcgttacatattgctggtatatcatcaattcttggagctattaattttattactacaattttaaatatacatcatataaactttactatagataaaatccctctcctagtatgatctatccttattacagctgtacttttattactatctctcccagtccttgctggagcaattactatactattaaccgatcgaaacttaaataccacttttttgatccaagtgggggggggatcctattttatatcaacatttattttgatttttggtcatcccgaagtttatattttaattttacctgga |
| J1D | tttgactcctacctcctcaatctccttattaattttaagaaacttcattggaagaggtgttgggactggttgaactatttatccccctcttgcttcaaacatttttcacagaggtccttcaattgatatatctatttctcgttacatattgctggtatatcatcaattcttggggctattaattttattactacaattttaaatatacatcatataaactttactatagataaaatccctctcctagtatgatctatccttattacagctgtacttttattactatctctcccagtccttgctggagcaattactatactattaaccgatcgaaacttaaataccacttttttgatccaagtgggggaggggatcctattttatatcaacatttattttgatttttggtcatcccgaagtttatattttaattacctgga |
| J1E | tttgactcctacctccatcaatctccttattaattttaagaaacttcattggaagaggtgttgggactggttgaactatttatccccctcttgcttcaaacatttttcacagaggtccttcaattgatatatctattttctcgttacatattgctggtatatcatcaattcttggagctattaattttattactacaattttaaatatacatcatataaactttactatagataaaatccctctcctagtatgatctatccttattacagctgtacttttattactatctctcccagtccttgctggggcaattactatactattaaccgatcgaaacttaaataccacttttttgatccaagtgggggaggggatcctattttatatcaacatttattttgatttttggtcatcccgaagtttatattttatttacctgg |

| | a |
|---|---|
| J2 | |
| J2A | ttttactcctcccatctccttattaattctaagaaactttattggaggcg<br>gtgttgggactggatgaaccatttatcctcctcttgcttcaaatatttt<br>ccatagaggtccctcagttgatatatctattttctcattacatattgct<br>ggtatatcatcaattcttggagctattaattttattactacaattttaaa<br>tatacatcacataaactttactatagataaaattcctctcttagtatg<br>atctatcctcattacagcagtacttttattattgtctctcccagttctcg<br>ccggagcaattactatactattaactgatcgaaacttaaatacca<br>ctttttttgatccaagaggagggggagaccctattttataccaacat<br>ttattttgattttttggccatcccgaagtttatattttatacctgga |
| J2B | tttgactcctccctccatcaatctccttattaattctaagaaactttatt<br>ggaggcggtgttgggactggatgaaccatttatcctcctcttgcttc<br>aaatattttccatagaggtccctcagttgatatatctattttctcattac<br>atattgctggtatatcatcaattcttggagctattaattttattactaca<br>attttaaatatacatcacataaactttactatagataaaattcctctct<br>tagtatgatctatcctcattacagcagtacttttattattgtctctccca<br>gttctcgccggagcaattactatactattaactgatcgaaacttaa<br>ataccactttttttgatccaaggggagggggagaccctattttatac<br>caacatttattttgattttttggccatcccgaagtttatattttaattttac<br>ctgga |
| J2C | tttgactcctccctccatcaatctccttattaattctaagaaactttatt<br>ggaggcggtgttgggactggatgaaccatttatcctcctcttgcttc<br>aaatattttccatagaggtccctcagttgatatatctattttctcattac<br>atattgctggtatatcatcaattcttggagctattaattttattactaca<br>attttaaatatacatcacataaactttactatagataaaattcctctct<br>tagtatgatctatcctcattacagcagtacttttattattgtctctccca<br>gttctcgccggagcaattactatactattaactgatcgaaacttaa<br>ataccactttttttgatccaggggggagggggagaccctattttatac<br>caacatttattttgattttttggccatcccgaagtttatattttaattttac<br>ctgga |
| J2D | tttgactcctccctccatcaatctccttattaattctaagaaactttatt<br>ggaggcggtgttgggactggatgaaccatttatcctcctcttgcttc<br>aaatattttccatagaggtccctcagttgatatatctattttctcattac<br>atattgctggtatatcatcaattcttggagctattaattttattactaca<br>attttaaatatacatcacataaactttactatagataaaattcctctct<br>tagtatgatctatcctcattacagcagtacttttattattgtctctccca<br>gttctcgccggagcaattactatactattaactgatcgaaacttaa<br>ataccactttttttgatccaaggggaggaggagaccctattttatac<br>caacatttattttgattttttggccatcccgaagtttatattttaattttac<br>ctgga |
| J2E | tttgactcctccctccatcaatctccttattaattctaagaaactttatt<br>ggaggcggtgttgggactggatgaaccatttatcctcctcttgcttc<br>aaatattttccatagaggtccctcagttgatatatctattttctcattac<br>atattgctggtatatcatcaattcttggagctattaattttattactaca<br>attttaaacatacatcacataaactttactatagataaaattcctctc<br>ttagtatgatctatcctcattacagcagtacttttattattgtctctccca<br>gttctcgccggagcaattactatactattaactgatcgaaacttaa<br>ataccactttttttgatccaagaggagggggagaccctattttatac<br>caacatttattttgattttttggccatcccgaagtttatattttaatttttac<br>ctgga |
| J2F | tttgactcctccctccatcaatctccttattaattctaagaaactttatt |

| | |
|---|---|
| | ggaggcggtgttgggactggatgaaccatttatcctcctcttgcttc<br>aaatattttccatagaggtccctcagttgatatatctattttctcattac<br>atattgctggtatatcatcaattcttggggctattaattttattactaca<br>attttaaatatacatcacataaactttactatagataaaattcctctct<br>tagtatgatctatcctcattacagcagtacttttattattgtctctccca<br>gttctcgccggagcaattactatactattaactgatcgaaacttaa<br>ataccactttttttgatccaggggggaggggggagaccctattttatac<br>caacatttattttgattttttggccatcccgaagtttatattttaatttttac<br>ctgga |
