## Supplementary Figures for "Shifts in demography in changing ecological conditions in a dependent-lineage population of harvester ant colonies"

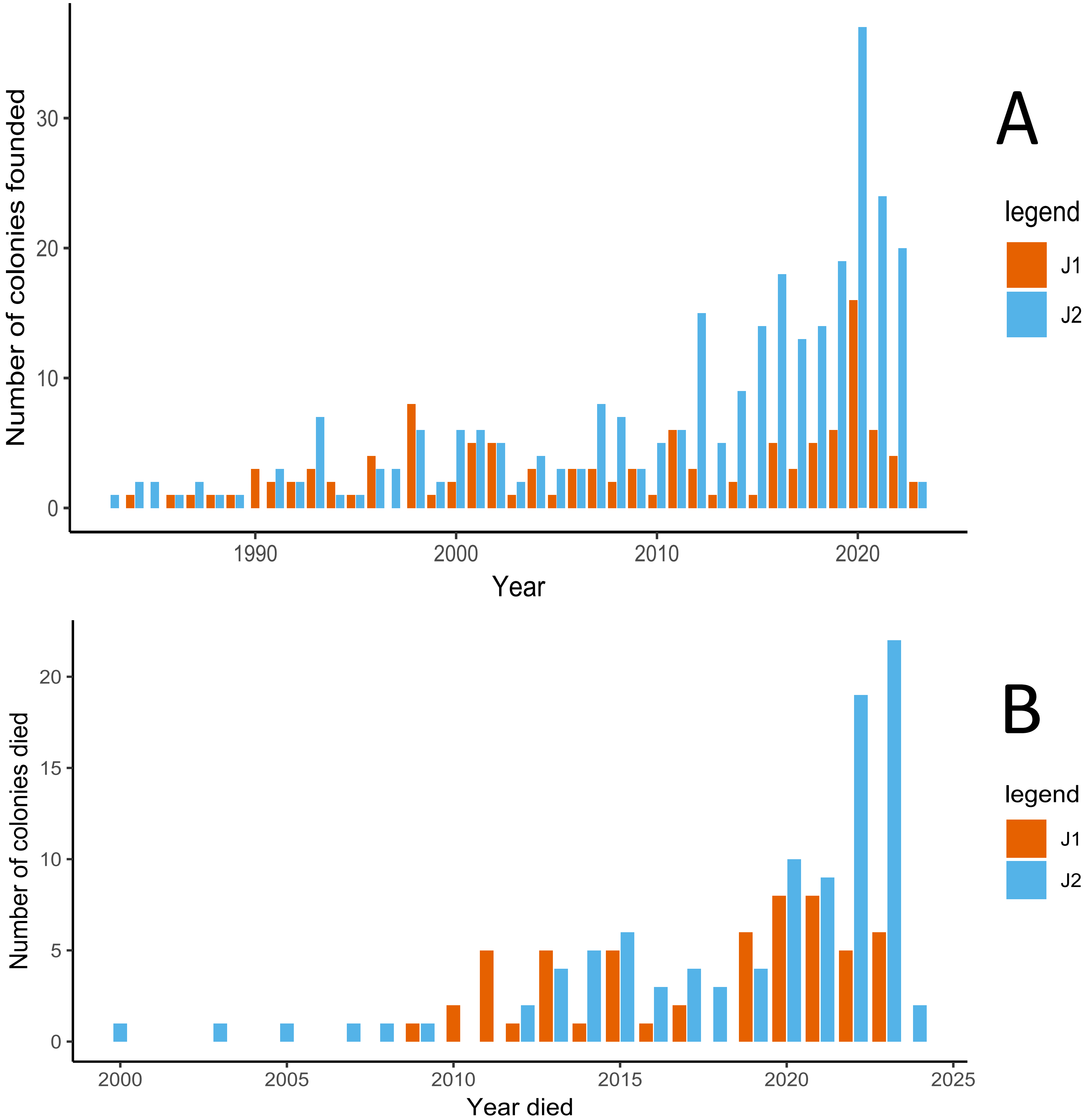


Supplementary Figure 1: Number of colonies in the sample set found per year (A) and number of colonies in the sample set that died per year (B)

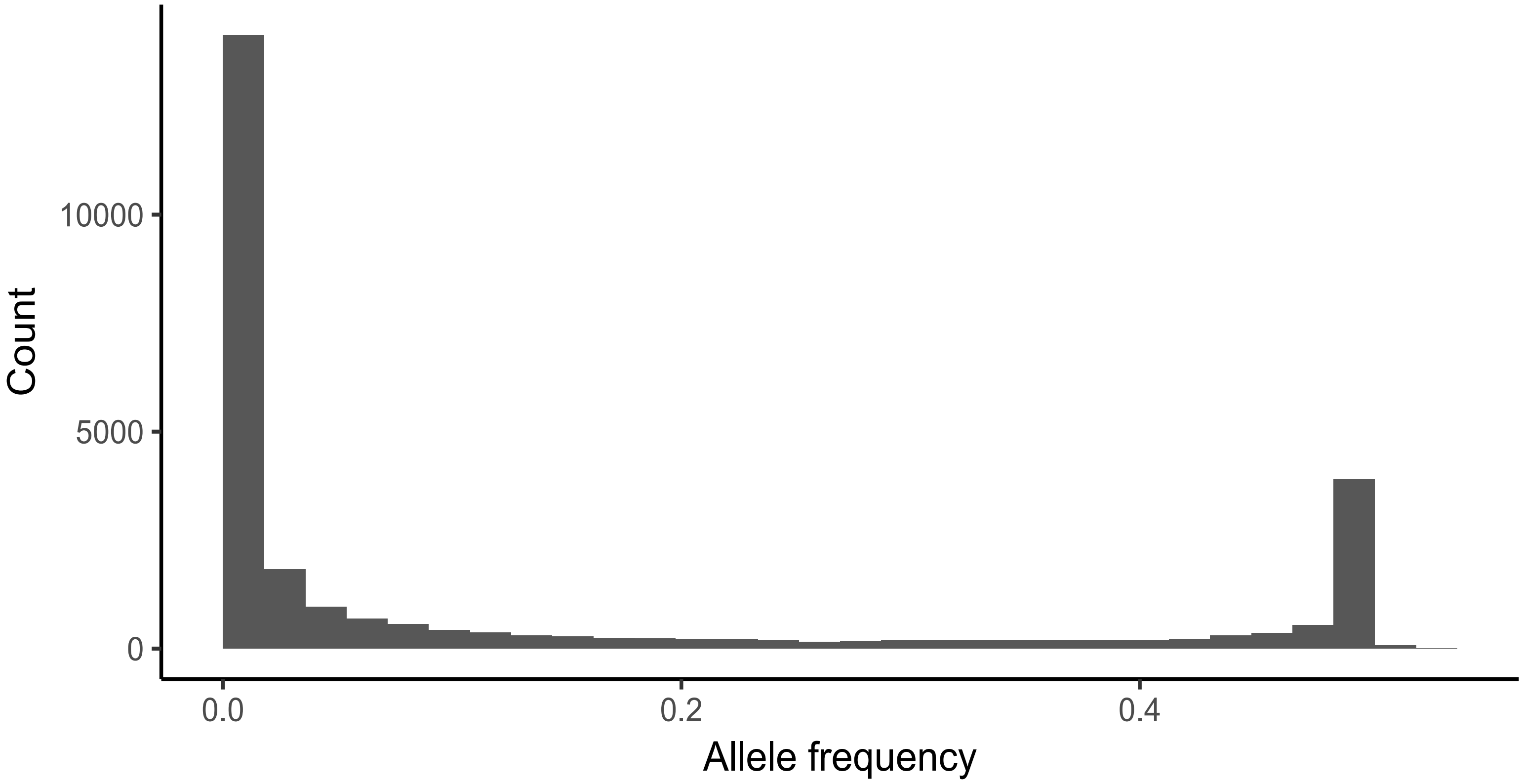


Supplementary Figure 2: Count of heterozygous genotype per locus
